# Microarray screening reveals a non-conventional SUMO-binding mode linked to DNA repair by non-homologous end-joining

**DOI:** 10.1101/2021.01.20.427433

**Authors:** Maria Jose Cabello-Lobato, Matthew Jenner, Christian M. Loch, Stephen P. Jackson, Qian Wu, Matthew J. Cliff, Christine K. Schmidt

## Abstract

SUMOylation is critical for a plethora of cellular signalling pathways including the repair of DNA double-strand breaks (DSBs). If misrepaired, DSBs can lead to cancer, neurodegeneration, immunodeficiency and premature ageing. Based on systematic proteome microarray screening combined with widely applicable carbene footprinting and high-resolution structural profiling, we define two non-conventional SUMO2-binding modules on XRCC4, a DNA repair protein important for DSB repair by non-homologous end-joining (NHEJ). Mechanistically, interaction of SUMO2 with XRCC4 is incompatible with XRCC4 binding to at least two other NHEJ proteins – XLF and DNA ligase 4 (LIG4). These findings are consistent with SUMO2 interactions of XRCC4 acting as backup pathways at different stages of NHEJ, in the absence of these factors or their dysfunctioning. Such scenarios are not only relevant for carcinogenesis, but also for the design of precision anti-cancer medicines and the optimisation of CRISPR/Cas9-based gene editing. This work reveals insights into topology-specific SUMO recognition and its potential for modulating DSB repair by NHEJ. Moreover, it provides a rich resource on binary SUMO receptors that can be exploited for uncovering regulatory layers in a wide array of cellular processes.

## Introduction

Posttranslational modification (PTM) with SUMO (small ubiquitin-like modifier) is key to regulating a gamut of cellular signalling pathways, including transcription, chromatin organisation, nuclear trafficking, DNA replication and DNA repair^1–5^. It is therefore not surprising that deregulation of the SUMO system is associated with a range of prevalent human diseases including neurodegenerative disorders, cardiovascular diseases and cancer^6–8^. In humans, several SUMO paralogues exist, with SUMO1-3 being ubiquitously expressed and established as posttranslational modifiers. SUMO2 and SUMO3 are almost identical, sharing 97% sequence identity. A lower sequence identity of ~45% between SUMO1 and SUMO2/3 results in more pronounced differences e.g. in their electrostatic surface potential. SUMOylation is mediated by an enzymatic triad, consisting of an activating E1 enzyme – a heterodimer formed by UBA2 (aka SAE2) and SAE1, the conjugating E2 enzyme UBE2I (aka UBC9), and one of ~10 E3 ligases. SUMOylation can occur on one or multiple lysines of substrate proteins, and as monomers or chains of multiple SUMO moieties, creating a complex array of topologies termed the SUMO code. PolySUMO chains in cells are primarily formed by SUMO2/3 linked via their internal K11 residues, while SUMO1 is mainly deemed a chain terminator. In analogy to the distinct functions assigned to different ubiquitin topologies, biochemical outcomes for distinct SUMO architectures can differ. Indeed, polySUMO chains are formed particularly in response to different types of stress, suggesting their importance in stress-related pathways. Despite our vast knowledge of ubiquitin chain functions, we still know very little about how polySUMO chains regulate specific cell signalling events^9–11^. By translating SUMOylations into defined biochemical actions, SUMO receptors – proteins non-covalently binding and recognising SUMO topologies – play a key role in determining the functional outcomes of SUMOylation events. Despite their importance and the large number (>7,000) of substrate SUMOylations existing in human cells^12^, only few (several tens) of SUMO receptors have been validated, and even less have been characterised for their binding to different SUMO topologies^13^. As a consequence, little is known about length- and paralogue-specific recognition of SUMO topologies. Indeed, in contrast to ubiquitin receptors, knowledge of different SUMO-binding modes is mainly limited to a small number of varying themes centred on 4-5 hydrophobic amino acids called SUMO interacting motifs (SIMs)^13–19^. These knowledge gaps limit our understanding of how SUMO functions at a mechanistic level and how it can best be exploited for treating human diseases associated with SUMO dysfunction and other purposes.

Here, we systematically screen the human proteome for receptors of polySUMO2 chains, identifying hundreds of candidates with diverse roles in established and emerging areas of SUMO biology. We validate a substantial and functionally varied set of SUMO receptors ensued by in-depth characterisation of the SUMO-binding modules of one of the identified receptors, XRCC4. XRCC4 is a core DNA repair factor known for its importance in DNA double-strand break (DSB) repair by non-homologous end-joining (NHEJ)^20^. DNA damage occurs frequently and can be caused by endogenous and exogenous sources. DSBs are the most cytotoxic DNA lesions and if left mis- or unrepaired, they can lead to cell death, mutagenesis or chromosomal translocation, and in turn cancer^21,22^. Cells have evolved two major pathways to repair DSBs: the first, homologous recombination (HR), repairs DSBs with high fidelity in late S/G2 cell cycle phases using a homologous sequence as a template, usually the sister chromatid; the second, NHEJ, is less accurate than HR, functions throughout interphase and is responsible for repairing the vast majority of DSBs in mammalian cells^21,22^. The importance for, and underlying mechanisms of, the SUMO system for key aspects of DSB repair by HR are well established, with SUMOylations of various HR factors and their decoding mechanisms via downstream receptors characterised^23,24^. By contrast, little is known about how SUMOylation regulates NHEJ, and no SUMO receptor roles have been defined for core NHEJ factors to date^24^. Here, we identify and characterise two distinct non-conventional polySUMO2-binding modules on XRCC4 located in its head and coiled-coil domains. Due to their location with respect to its known DNA repair-important regions, XRCC4 binding to SUMO2 has potential to regulate NHEJ at distinct stages linked to the functions of at least two other core NHEJ factors, XLF and LIG4.

## Results

### Proteome microarray screening retrieves polySUMO2 receptors enriched for diverse gene ontologies

SUMO receptors have mainly been identified using protein-protein interaction (PPI) screens based on yeast-two-hybrid systems, and affinity purification from whole cell extracts using SUMO topologies as baits combined with mass spectrometry^16,25–29^. These technologies are limited by their propensity to identify indirect SUMO binders in addition to direct, binary SUMO receptors. Moreover, yeast-two-hybrid systems are restricted to gene-encoded linear fusion topologies rather than the enzymatically generated SUMO chains existing in cells. Additionally, mass spectrometry-based approaches are limited to the cell- and tissue-specific proteomes used as starting materials and display a bias towards abundant proteins. To overcome these limitations, we systematically screened the human proteome for polySUMO2 receptors using microarrays containing duplicate protein spots of ~15,000 unique full-length human genes. To this end, we hybridised the arrays with enzymatically linked polySUMO2 chains fluorescently labelled with Cy5, followed by fluorescence scanning and background subtraction using soluble Cy5 as a reference (Fig. 1A).

**Figure 1.**
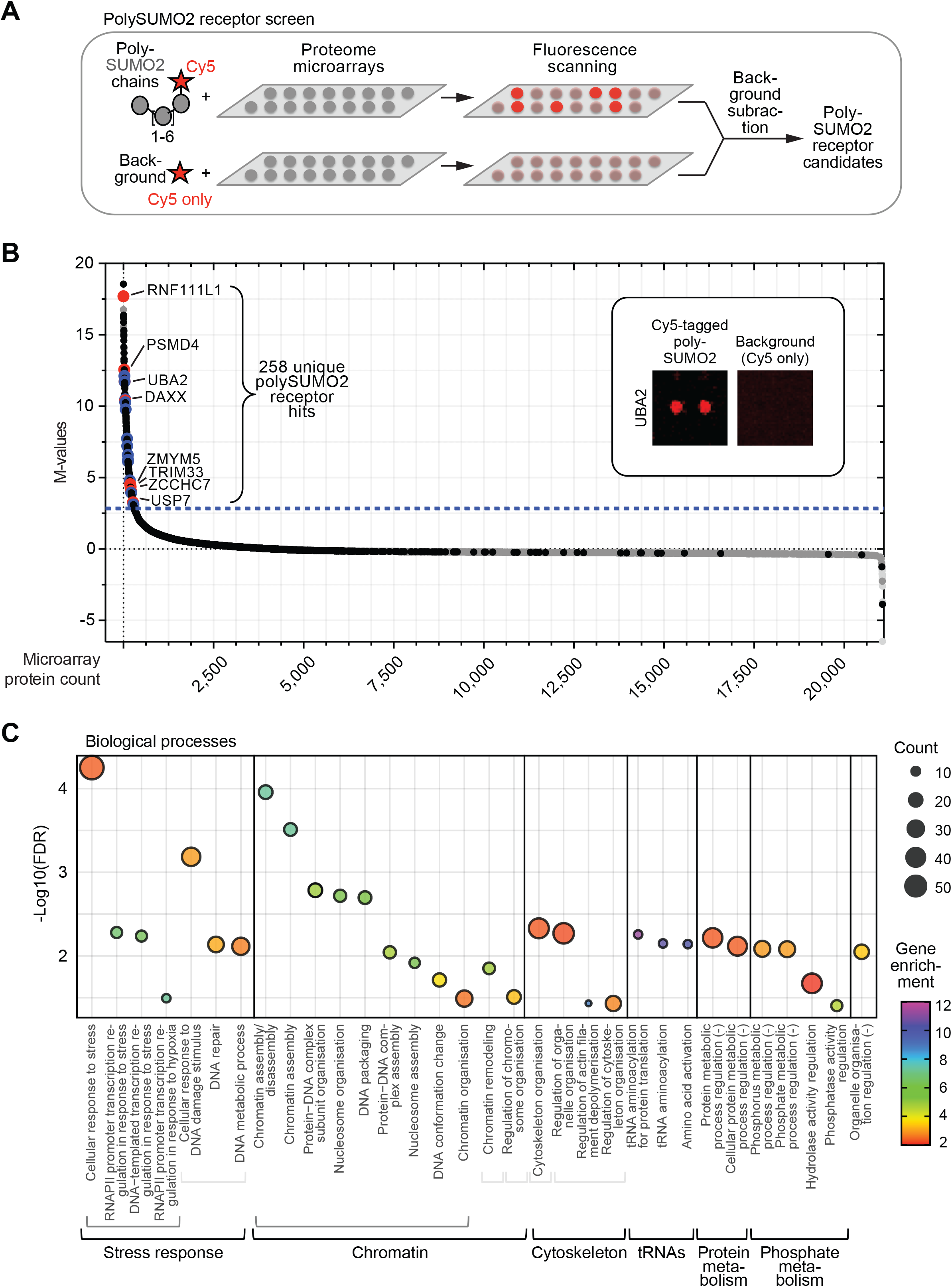
Protein microarray screen identifies novel polySUMO2 receptors. (**A**) Human proteome microarray screening pipeline. (**B**) Screening results highlighting known/described SUMO2 receptors (red) and ones validated in this study (blue). Threshold for polySUMO2 receptor candidates (dashed blue line) set to 2.85. For details see Methods section. (**C**) Gene ontology enrichment for biological processes enriched >2-fold in SUMO receptor candidate list, using Panther Classification System^129,130^. (-) denotes negative regulation. FDR: false discovery rate; RNAPII: RNA polymerase II.

The screen identified a total of 258 unique binary polySUMO2 receptor hits (Fig. 1B; Supplementary Tables S1, S2), featuring known SUMO2 receptors as well as a large number of proteins with no previous SUMO-binding functions assigned to them. As expected, known SUMO receptors harboured components of the SUMO conjugation cascade, in addition to downstream receptors with no known SUMOylation roles. As such, SUMO receptors/SUMOylation components amongst the hits included the ubiquitin E3 ligase, RNF111L1, also known as Arkadia-like 1 (ARKL1)^16^, one of the top two hits; UBA2, a component of the heterodimeric SUMO E1 enzyme^30–33^; the chromatin regulator DAXX^34–36^ – one of the best characterised SUMO receptors; the zinc finger proteins ZMYM5 (also known as ZNF237) and ZCCHC7^16,37^; TRIM33 – a ring finger protein that can act as a SUMO E3 ligase^38^; and the deubiquitylase USP7 (also known as HAUSP)^39^. In addition, PSMD4 (also known as S5A) was amongst the hits, a non-ATPase regulatory 26S proteasome subunit implicated as a potential receptor for hybrid SUMO-ubiquitin chains^40^. These data highlight the feasability of our approach to identify binary SUMO receptors. The approach is unlimited by the receptors binding to a specific region on SUMO, thereby overcoming restrictions of other recently developed, complementary methodologies^41^. Moreover, as expected, SUMO receptors with known preferences for SUMO1 binding e.g. RGS17^42^, DPP9^43^, and PARK2^44^, did not score as hits, suggesting our approach was able to distinguish between different SUMO paralogues. Known SUMO receptors tend to be large proteins with an average length of >700 amino acids, compared to an average human protein length of ~400 amino acids^45^. Indeed, a substantial number of SUMO receptors are more than 1,500 amino acids long, including HERC2^46^, RANBP2 (also known as NUP358)^19^, CENPE^47^, SETX^16^, P300^48^, CHD3^49^, CASP8AP2 (also known as FLASH)^17^ and SLX4^50^. Due to the increased challenge of purifying large proteins, microarrays likely feature smaller proteins with increased structural functionality and/or lack proteins with particularly high molecular weights, explaining the absence of some of these receptors from the microarrays and/or the candidate list (Fig. 1B; Supplementary Tables S1, S2). Other SUMO receptors missing from the list may interact more efficiently with SUMO architectures not employed in our screen.

SUMOylation is especially known for its importance in regulating processes inside the nucleus and in response to stress^51,52^. Consistent with this notion, gene ontology analysis of the polySUMO2 hits resulted in a significant enrichment of biological processes linked to stress-induced transcription and DNA repair, in particular in response to hypoxia and DNA damage^24,51–54^ (Fig. 1C). These findings underpin the validity of our approach in identifying receptors in pathways associated with known SUMO functions, and indeed in contexts triggering the formation of polySUMO chains, an achievement previously unattained with proteome microarrays^55^. Other significantly enriched ontologies included chromatin-associated processes such as remodelling and nucleosome assembly/disassembly that are also linked to SUMO function^3,56^ (Fig. 1C). Notably, significant enrichment was also observed for cytoplasmic processes, such as cytoskeletal organisation, consistent with SUMOylation emerging as an important regulatory layer in that area^57,58^. tRNA aminoacylation, a field less established for regulation by SUMOylation featured amongst the most strongly enriched biological processes. Finally, our analyses connected SUMO processes to phosphorylation. In this regard it is noteworthy that cross-talk between different PTMs represents an emerging and exciting theme in the ubiquitin and ubiquitin-like protein fields^59,60^. Strikingly, the majority of the receptor candidates were proteins uncharacterised for SUMO-binding functions independently of whether they were assigned to expected gene ontologies or to emerging SUMO functions. Overall, these findings highlight our screening platform as a feasible approach for revealing known and hitherto unidentified SUMO receptors involved in a vast array of cell biology areas with established as well as understudied connections to known SUMO biology aspects.

### Biolayer interferometry validates polySUMO2 receptors with diverse functions and binding characteristics

Having identified a range of biological processes significantly enriched amongst the hits, we next addressed if any of the candidates were functionally interconnected. To this end, a STRING network analysis revealed a significant enrichment of PPIs amongst the hits (enrichment *p*-value = 1.51×10^−12^). Specifically, the analysis elucidated several gene clusters connected via a central hub of processes associated with p53 function (Fig. 2A). Similarly to the enriched ontologies (Fig. 1C), some, but not all, of these clusters were linked to SUMO-associated functions, as demonstrated by the presence of known SUMO receptors amongst some of them (Fig. 2A). For example, the coordinated functions of DAXX and USP7 to regulate p53 function^61^ connected them to the central p53-associated cluster, and PSMD4 was linked to several other proteasomal components not previously associated with SUMO binding. In this regard, both VCP (also known as p97) and one of its interaction partners, NSFL1C, came up as hits. Notably, a different interaction partner of VCP, UFD1, functions as a SUMO receptor in yeast to help recruit the VCP complex to its targets^62^. This finding raises the possibility that the VCP complex could take on similar roles in humans. Other gene clusters formed by SUMO receptor candidates centred on nuclear functions including DNA repair and chromatin-associated processes as well as pre-mRNA splicing^63,64^, consistent with enrichment of these pathways also in our gene ontology analysis (Fig. 1C). Interestingly, several serine/threonine kinases (p21-activating kinases; PAKs) formed part of a gene cluster linking cytoplasmic functions associated with the cytoskeleton to nuclear signalling. These factors have not directly been associated with SUMO functions. Finally, five aminoacyl tRNA synthetases formed a separate cluster, representative of the strong enrichment of tRNA-associated processes in the identified gene ontologies (Fig. 1C). None of these proteins had previous SUMO receptor functions assigned to them. tRNA transcription is regulated by SUMOylation in response to stress in yeast^65,66^, raising the possibility that SUMOylation, and SUMO receptor functions, in tRNA-mediated protein translation could help cells respond to stress^65^, thereby offering a starting point to assess such functions mechanistically.

**Figure 2.**
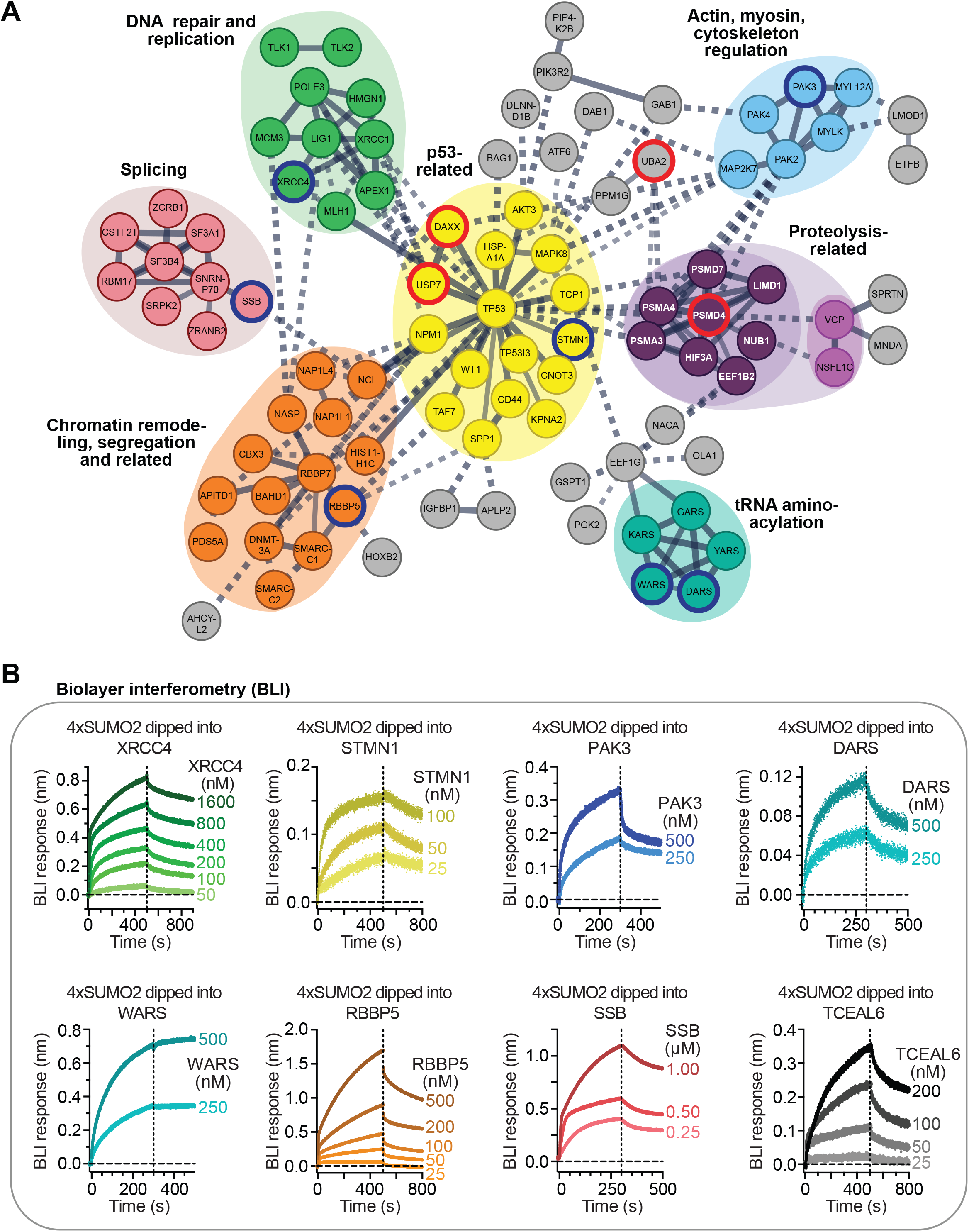
Network analysis and validation of polySUMO2 receptor candidates. (**A**) Clusters of polySUMO2 receptor candidates identified by STRING network analysis^131^. Red and blue strokes highlight known SUMO receptors, and receptors validated in this study, respectively. (**B**) Validation of polySUMO2 chain receptor candidates across different gene clusters and with no consensus SUMO-interacting motifs (SIMs) present in their sequence using biolayer interferometry (BLI). Colouring according to corresponding gene clusters in (A). Association and dissociation phases are separated by vertical dashed lines, as indicated.

To assess the validity of the identified receptor candidates, we performed SUMO-binding assays of a range of candidates using biolayer interferometry (BLI), a biophysical technique related to surface plasmon resonance (SPR) (Fig. 2B; Supplementary Fig. S1). Using genetically encoded linear SUMO2 chains, fused via their C-terminal diglycine and lysine 11 of the distal and proximal moieties (4xSUMO2), a well established model topology^67–71^, we validated eight candidates distributed across a range of gene clusters and M-values (Fig. 1B; Supplementary Table S1). These included the DNA repair protein XRCC4; STMN1 – a cytosolic protein regulated by p53 that has recently been hypothesised to exert its function via SUMO binding^72^; the serine/threonine protein kinase PAK3; two aminoacyl-tRNA synthetases – DARS and WARS; RBBP5 – a transcriptional regulator associated with histone methyltransferase complexes, and SBB – a protein important for various aspects of RNA metabolism (Fig. 2B, genes with blue frames). We also validated the transcriptional elongation factor TCEAL6 for which no putative SIMs could be retrieved using JASSA and GPS-SUMO, two state-of-the-art SIM prediction servers^13,73^. Notably, the eight validated SUMO2 receptors displayed a range of association and dissociation profiles, with some associating notably more stably with 4xSUMO2 than others (Fig. 2B, compare e.g. WARS to TCEAL6). The presence of at least one SUMO2 receptor, previously known or validated in this study, in every identified gene cluster emphasises the overall validity of our screening approach. Taken together, these results demonstrate the ability of our screen to identify SUMO2 receptors with distinct binding characteristics, featuring a wide range of biological functions and highlighting established as well as hitherto undiscovered links to SUMO binding and functionality.

### XRCC4 preferentially binds to polySUMO2/3 chains over shorter topologies

To further define the SUMO-binding characteristics of one of the receptors arising from our screen, we selected the core NHEJ factor XRCC4 (M-values ~6; Fig. 1B and Supplementary Table S1) for follow-on studies. SUMOylation is known to be crucial for efficient NHEJ to take place^74–76^. Despite this importance, knowledge on SUMO receptor roles for core NHEJ factors is lacking. XRCC4 is a 38 kDa protein that exists mainly as a homodimer in cells and features a number of functionally and structurally distinct domains^77–79^. Its N-terminal head domain is important for interaction with another NHEJ core factor, XLF, followed by a coiled-coil domain that mediates binding with the NHEJ ligating enzyme LIG4 or the nucleoskeleton protein IFFO1^78,80–83^. A flexible C-terminal tail is important for interacting with other NHEJ-associated DNA repair factors^84–87^ (Fig. 3A). NHEJ is initiated in response to DSBs, with Ku, a heterodimer formed by Ku70 and Ku80, recognising and binding to broken DNA ends. Amongst several functions, DNA-bound Ku acts as a recruitment hub for other NHEJ factors e.g. the DNA damage response kinase DNA-PKcs, which together with Ku forms the holoenzyme DNA-PK (Fig. 3B). DNA-PK itself can recruit other factors important for preparing the broken DNA ends for ligation^88,89^. XRCC4 is key for facilitating distinct aspects of NHEJ. By interacting with the core NHEJ factor XLF, XRCC4 is implicated in tethering DNA ends and thus, promoting synapsis and subsequent DNA-end ligation, with the importance of this varying depending on cellular context^90–95^. Moreover, XRCC4 interaction with IFFO1 contributes to spatial stabilisation of the broken ends to prevent chromosomal translocations^81^. A separate population of XRCC4 interacts with LIG4 instead of IFFO1. In addition to stabilising LIG4 and promoting its enzymatic activity, XRCC4 helps recruit LIG4 to broken DNA ends, thereby facilitating the final ligation step necessary for NHEJ to conclude (Fig. 3B)^96^.

**Figure 3.**
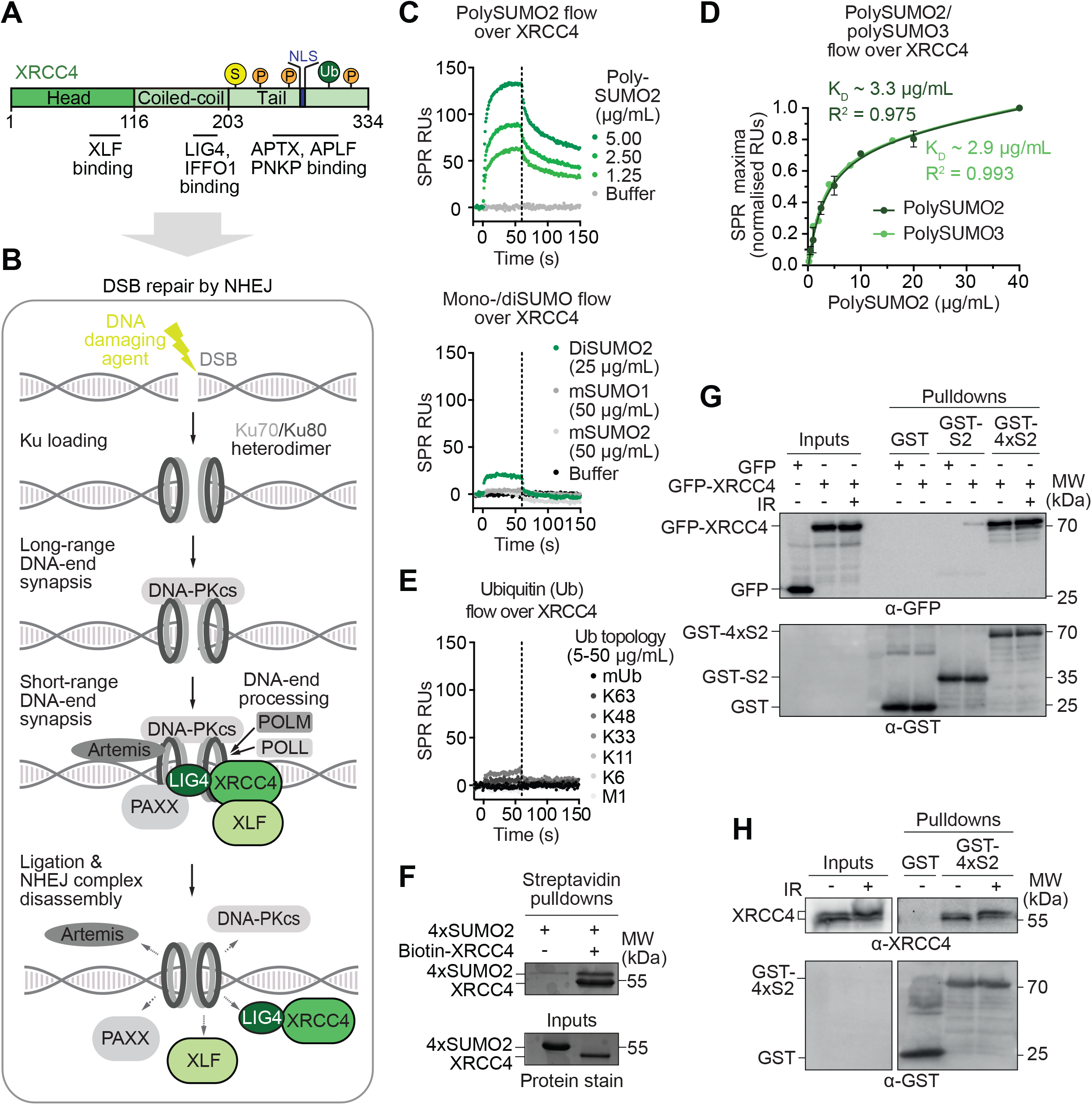
XRCC4 preferentially binds to polySUMO2 chains. (**A**) Protein schematic of XRCC4 highlighting key structural and functional features. S: SUMOylation; P: phosphorylation; Ub: ubiquitylation; NLS: nuclear localisation signal. (**B**) Schematic of DNA double-strand break (DSB) repair by non-homologous end-joining (NHEJ). (**C**) SPR sensorgrams showing preferential binding of XRCC4 to polySUMO2 chains (top) over SUMO monomers (mSUMO1, mSUMO2) and dimers (diSUMO2; bottom), using XRCC4 as ligand and SUMO topologies as analytes. Association and dissociation phases are separated by vertical dashed lines, as indicated. (**D**) Equilibrium analysis of SPR response unit maxima of polySUMO2 binding to immobilised wildtype (WT) and mutant versions of full-length XRCC4, showing binding of XRCC4 to polySUMO2 and polySUMO3 chains with similar affinity. (**E**) SPR sensorgram showing no or minor detectable binding of XRCC4 to ubiquitin monomers (mUb) and tetra-ubiquitin chains linked via ubiquitin’s internal lysines K63, K48, K33, K11, K6 or its N-terminal methionine (M1), using XRCC4 as ligand and ubiquitin topologies as analytes. Association and dissociation phases are separated by vertical dashed lines, as indicated. (**F**) XRCC4 interacts with 4xSUMO2 (6xHis-4xSUMO2-Strep) in solution in vitro. (**G**) XRCC4 preferentially associates with SUMO2 chains over SUMO2 monomers in solution as shown by GST-mSUMO2 and GST-4xSUMO2 pulldowns co-precipitating GFP-XRCC4 ectopically expressed in HEK293T XRCC4 −/− cells in the absence or presence of DNA damage induced by ionizing radiation (IR; 15 Gy, ~15 min). Inputs were 4% of the total. (**H**) GST-4xSUMO2 pulldowns co-precipitate endogenous XRCC4 from HEK293T nuclear extracts in the absence or presence of IR-induced DNA damage (15 Gy, ~15 min). K_D_: dissociation constant; RUs: response units; S2: SUMO2; SPR: surface plasmon resonance.

Given that different topologies of SUMO are associated with distinct functions^2,52,97^, we first assessed if XRCC4 displayed preferential binding to certain SUMO topologies. Indeed, SPR assays revealed selective binding of XRCC4 to enzymatically linked polySUMO2 chains known to exist in cells^10,98–100^ (Fig. 3C, top) and consistent with our polySUMO2 microarray screen. Moreover, XRCC4 bound polySUMO2 chains preferentially over SUMO1/2 monomers (mSUMO1/2) and SUMO2 dimers (diSUMO2; Fig. 3C, bottom; Supplementary Fig. S2A). In agreement with the high sequence identity between SUMO2 and SUMO3, XRCC4 bound to polySUMO2 and polySUMO3 chains with similar affinity (dissociation constant K_D_ ~ 3 μg/mL; Fig. 3D; Supplementary Fig. S2B). Taken together, these findings suggested the existence of multiple SIMs on XRCC4 that bind to distinct SUMO moieties of the same chain with increased avidity.

Since SUMO belongs to the ubiquitin/UBL family, we next investigated if XRCC4 showed a preference for SUMO-binding over other ubiquitin/UBL members. Using the same SPR setup as before, we detected no interaction of XRCC4 to a wide range of ubiquitin topologies (Fig. 3E; Supplementary Fig. S2C). Next, we performed precipitations with 4xSUMO2 as bait, demonstrating that recombinant XRCC4 was able to bind to SUMO2 chains not only on solid surfaces but also in solution (Fig. 3F). XRCC4 was also able to bind to SUMO2 in cells. Similarly to the binding characteristics we established in vitro, ectopically expressed XRCC4 preferentially co-precipitated from cellular extracts with polySUMO2 chains over SUMO2 monomers (mSUMO2), irrespectively of the presence or absence of ionizing radiation (IR)-induced DNA damage (Fig. 3G). Importantly, SUMO2 chains were also able to precipitate endogenous XRCC4 from cellular extracts, including high-molecular weight forms, representative of DNA damage-induced phosphorylation^101^ (Fig. 3H). The functions of SUMO topologies tend to be exerted via conjugation to substrates. To test if XRCC4 could bind to SUMO chains when covalently bound to another protein, we performed pulldown assays with SUMO2 fused to a model substrate (tetracycline repressor), which readily co-precipitated with XRCC4 (Supplementary Fig. S2D). Overall, these findings demonstrated that a population of XRCC4 in cells was able to, and available for, binding to polySUMO2 chains in the absence or presence of DNA damage and independently of whether SUMO existed as a free entity or was covalently bound.

### XRCC4 lacks functional consensus SIMs

To characterise if and how the positioning of SUMO-binding regions on XRCC4 related to the structure and function of XRCC4’s known domains (Fig. 3A), we next investigated if XRCC4 contained any conventional SIM sequences. SIMs feature a core of 4-5 hydrophobic residues that can be intersected or framed by negatively charged residues on one or both sides (Supplementary Fig. S3B)^18,16,19,28,102^. Indeed, JASSA^13^ and GPS-SUMO^73^ predicted five putative SIMs (pSIMs) on XRCC4: three located in its head domain (pSIMs 8, 33, and 123), one in its coiled-coil region (pSIM181), and one in the C-terminal part of the protein (pSIM257; Supplementary Fig. S3A). However, the crystal structures of XRCC4^77,78,82^ suggested that pSIMs 8, 33, and 123 are important for the structural integrity of the XRCC4 head domain by forming extensive interactions with nearby XRCC4 residues. Moreover, pSIM33 is almost completely buried inside the XRCC4 head domain, rendering this motif inaccessible to surface interactions^77^ (Supplementary Fig. S3A, note the lack of green in the structure on the right). Consistent with these realisations, alanine mutations of pSIMs 8, 33, or 123 abolished XRCC4 interactions not only with polySUMO2/3 (Supplementary Fig. S3C, D), but also with XLF (Supplementary Fig. S3E, F), which binds to a region on XRCC4’s head domain well separated from the pSIM locations^83,90,103,104^ (compare Supplementary Fig. S3A to Fig. 3A). By contrast, alanine mutation of pSIM181 neither affected polySUMO2/3 nor XLF binding of XRCC4 (Supplementary Fig. S3C-F). In addition, deletion of XRCC4’s C-terminal tail containing pSIM257 did not markedly affect polySUMO2 binding of XRCC4. Collectively, these data eliminated the five pSIMs as functional SIMs on XRCC4 (Supplementary Fig. S3G). We conclude that XRCC4 must interact with SUMO2/3 via a non-conventional, hitherto unidentified, binding mode.

### Carbene footprinting determines distinct SUMO2-binding regions on XRCC4

In light of the absence of conventional SIMs, we used a recently developed chemical biology approach, known as carbene footprinting^105–107^, to comprehensively map SUMO2 interactions along XRCC4. In contrast to other high-resolution PPI methods, this technology requires little starting material and is unrestricted by the molecular weight of the targeted protein as well as the affinity of the interaction. Indeed, NMR spectra of full-length XRCC4 were characterised by severely attenuated signals, preventing detailed analyses. Carbene footprinting is based on differential labelling of the surfaces of interaction partners with a photo-activated diazirine-containing probe, in this case an aryldiazirine, yielding highly reactive carbene species that rapidly label the protein surface. Labelling of the protein-of-interest with the aryldiazirine probe individually and as a mixture, followed by enzymatic digestion combined with mass spectrometry, allows comparative quantification of the resulting peptide labelling levels with high accuracy and resolution defined by peptide length and labelling efficiency. Reduced peptide labelling in the presence of a binding partner, indicate the residues of the peptide as potential binding sites due to surface masking. In addition, unmasking of peptides labellings can occur, and both masking and unmasking can indicate interaction-induced conformational changes or rearrangements leading to a change in exposure of residues to solvent, and therefore labelling^105–107^ (Fig. 4A).

**Figure 4.**
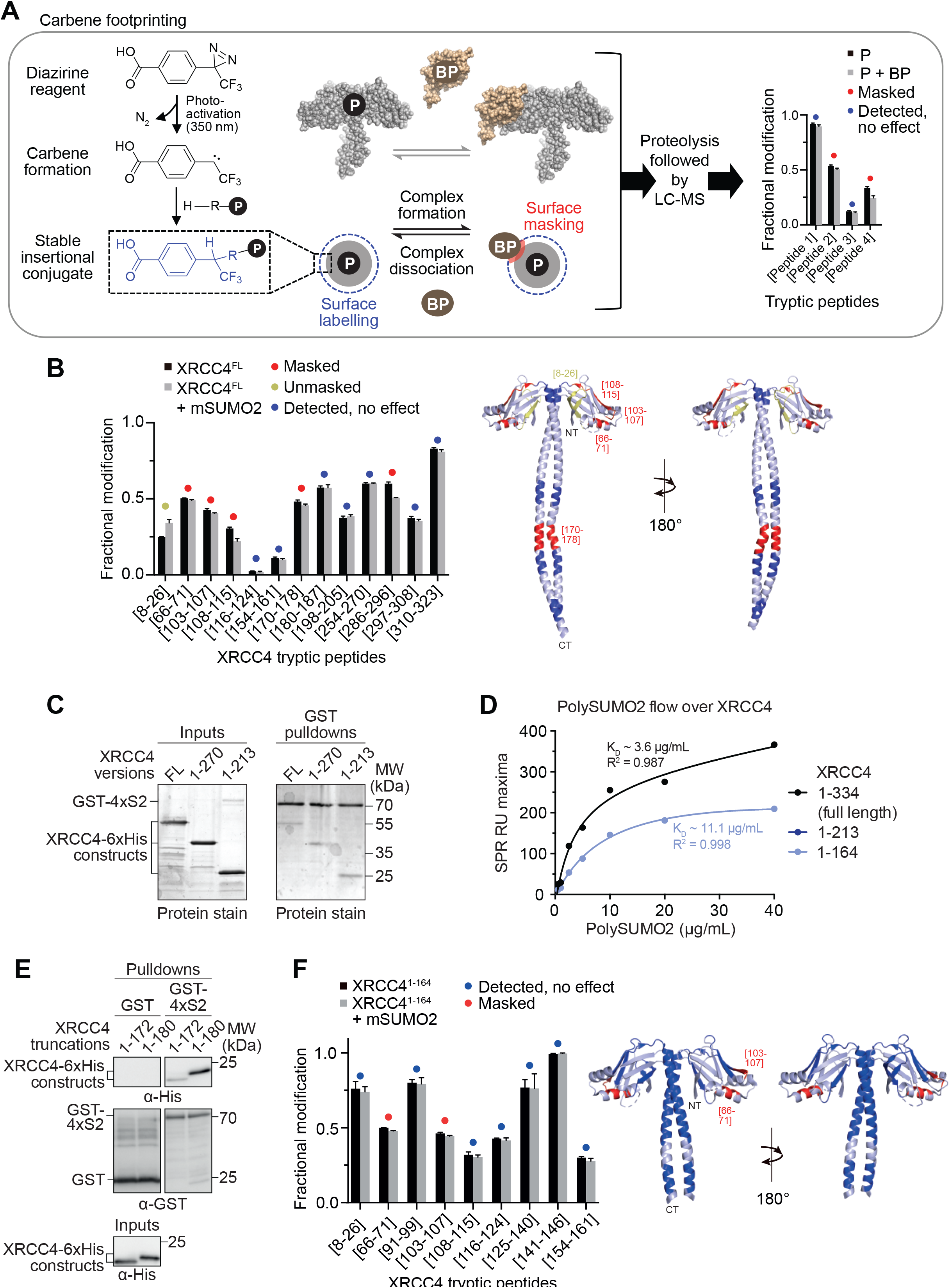
Carbene footprinting identifies distinct SUMO2-binding regions on XRCC4. (**A**) Experimental pipeline for carbene footprinting of the interaction between a protein (P)-of-interest, in this case XRCC4 (PDB 1IK9)^78^, and its binding partner (BP), in this case monomeric SUMO2 (mSUMO2; PDB 2N1W)^132^, based on chemical labelling of the XRCC4 surface using aryldiazirine as a photo-activated probe. Fractional labelling of XRCC4 with aryldiazirine, individually and in the presence of mSUMO2, followed by proteolysis (tryptic peptide digestion) combined with liquid chromatography-mass spectrometry (LC-MS), allows comparative quantification of the labelling levels of the digested peptides with high accuracy and resolution defined by peptide length and labelling efficiency. Any peptide labelling changes in the absence or presence of binding partner indicate changes in surface accessibility of the monitored peptide. XRCC4 and mSUMO2 (**B**) Fractional modification by aryldiazirine of full-length XRCC4 (XRCC4^FL^, representative protein stain shown in Supplementary Fig. S1) in the presence (grey bars) or absence (black bars) of SUMO2 monomers (mSUMO2). Error bars represent +/− standard deviations based on n=3 independent biological replicates. Significant peptide masking and unmasking differences (Student’s t-test, p<0.05) are highlighted in red or yellow, respectively, with unchanged peptide probing indicated in blue. The same colour code is applied to the structure of XRCC4^1-213^ (PDB 1IK9) on the right, with undetected regions indicated in light blue. (**C**) GST-4xSUMO2 pulldowns uncover comparable levels of recombinant full-length (FL) XRCC4, XRCC4^1-270^ and XRCC4^1-213^ in the precipitated fractions. (**D**) Steady-state affinity determination of polySUMO2 binding to immobilised XRCC4^FL^ (black), and XRCC4^1-164^ (blue) by surface plasmon resonance (SPR). (**E**) GST-4xSUMO2 pulldowns reveal higher levels of recombinant XRCC4^1-180^ in the precipitated fractions compared to XRCC4^1-172^. (**F**) Fractional modification by aryldiazirine of XRCC4^1-164^ in the presence (grey bars) or absence (black bars) of mSUMO2. Error bars represent +/− standard deviations based on n=3 independent biological replicates. Significant peptide masking and unmasking differences (Student’s t-test, p<0.05) are highlighted in red or yellow, respectively, with unchanged peptide probing indicated in blue. The same colour code is applied to the structure of XRCC4^1-164^ (PDB: 1IK9) on the right, with undetected regions indicated in light blue. K_D_: dissociation constant; CT: C-terminus; NT: N-terminus; RU: response unit; S2: SUMO2.

Carbene footprinting of XRCC4 in the presence of mSUMO2 revealed multiple XRCC4 peptides as potential SUMO2-interacting sites, consistent with the avidity we observed for polySUMO2/3 binding. Masking was detected in the XRCC4 head domain between residue positions 66-115, in the coiled-coil region (170-178 peptide), although the preceding peptide evaded labelling, and in the C-terminal tail in/around the 286-296 region. Unmasking occurred in the 8-26 region, an area on the head domain spatially proximal to the N-terminal part of the coiled-coil (Fig. 4B). XRCC4 truncations lacking up to 121 amino acids of the C-terminus were precipitated by GST-4xSUMO2 with similar levels compared to full-length XRCC4 (Fig. 4C), in agreement with the comparable polySUMO2 affinities we measured in SPR equilibrium analyses (Supplementary Fig. S3G). These findings suggested that the masking of the 286-296 region was due to structural rearrangements of XRCC4’s C-terminus rather than its direct involvement in SUMO binding. A 1-164 XRCC4 truncation (XRCC4^1-164^) retained substantial polySUMO2 binding, albeit with weaker affinity, as illustrated by a ~3-fold increase in its K_D_, consistent with XRCC4’s head domain contributing to SUMO binding (Fig. 4D). In addition to the head domain, increased precipitation of XRCC4^1-180^ by GST-4xSUMO2 compared to XRCC4^1-172^ confirmed the presence of residues important for SUMO binding in the coiled-coil (Fig. 4E). Altogether, these findings pointed towards SUMO-interacting regions on XRCC4 in its head and coiled-coil domains, with potential allosteric changes occurring at/around positions 8-26 and in the flexible C-terminus (286-296 positions). To increase the resolution of the carbene footprinting approach in the head domain, we repeated the analyses with XRCC4^1-164^, which retrieved an extended set of labelled peptides. Overall, the experiment consolidated the effects we observed with full-length XRCC4 (Fig. 4F), and strengthened the conclusion of at least two potential distinct SUMO-binding regions existing on the head domain localised to/around residue regions 66-71 and 103-107.

### XRCC4 head domain binds to SUMO2 in a non-conventional, paralogue-specific manner

Having narrowed down potential regions of SUMO binding to distinct and defined parts of XRCC4, we next performed NMR titrations of targeted XRCC4 truncations to map SUMO interactions at an increased – amino acid-level – resolution. To this end, we first established XRCC4^1-164^ as the largest head domain-containing XRCC4 construct amenable for NMR analysis. Two amino acid stretches showed marked intensity loss in the ^1^H-^15^N BEST-TROSY spectra. Addition of increasing concentrations of mSUMO2 resulted in differential attenuation of signal intensities, consistent with a specific interaction between mSUMO2 and XRCC4 with a discrete binding site. Residues 101-LKDVSFRLGSF-111 (henceforth referred to as SIM101) displayed the strongest effects, followed by residues 56-ADDMA-60 (henceforth termed SIM56), with both regions forming coherent and spatially proximal surface sites on XRCC4’s head domain (Fig. 5A; Supplementary Fig. S4A). Equivalent experiments with diSUMO2 and 4xSUMO2 highlighted the same amino acid stretches (Supplementary Figure S4B, C), further consolidating these regions as SUMO-binding surfaces on XRCC4. Additional affected residues likely reflect more extended contact regions due to the larger volumes occupied by the di- and 4xSUMO2 topologies compared to mSUMO2. Overall, these findings were in line with the results we obtained by carbene footprinting, highlighting peptide fractional modification as an invaluable technology for narrowing down SUMO-binding regions along full-length proteins, which can be a daunting task, particularly for large proteins lacking conventional SIMs. It is worth noting that none of the pSIMs predicted by JASSA and GPS-SUMO underwent comparable intensity losses (Supplementary Fig. S4A), consistent with our previous conclusion of XRCC4 lacking functional consensus SIMs. In contrast to mSUMO2, addition of mSUMO1 resulted in fewer, different and substantially less pronounced changes in the ^1^H-^15^N BEST-TROSY spectra of XRCC4^1-164^. In this regard we note that XRCC4 binding was undetectable in the ^1^H-^15^N HSQC spectra of mSUMO1 even after addition of 0.5 molar equivalents of XRCC4. Collectively, these data demonstrated selective binding of XRCC4 to SUMO2 over SUMO1 paralogues, and higher affinity for polySUMO2/3 chains over shorter SUMO topologies (Supplementary Figure S4D).

**Figure 5.**
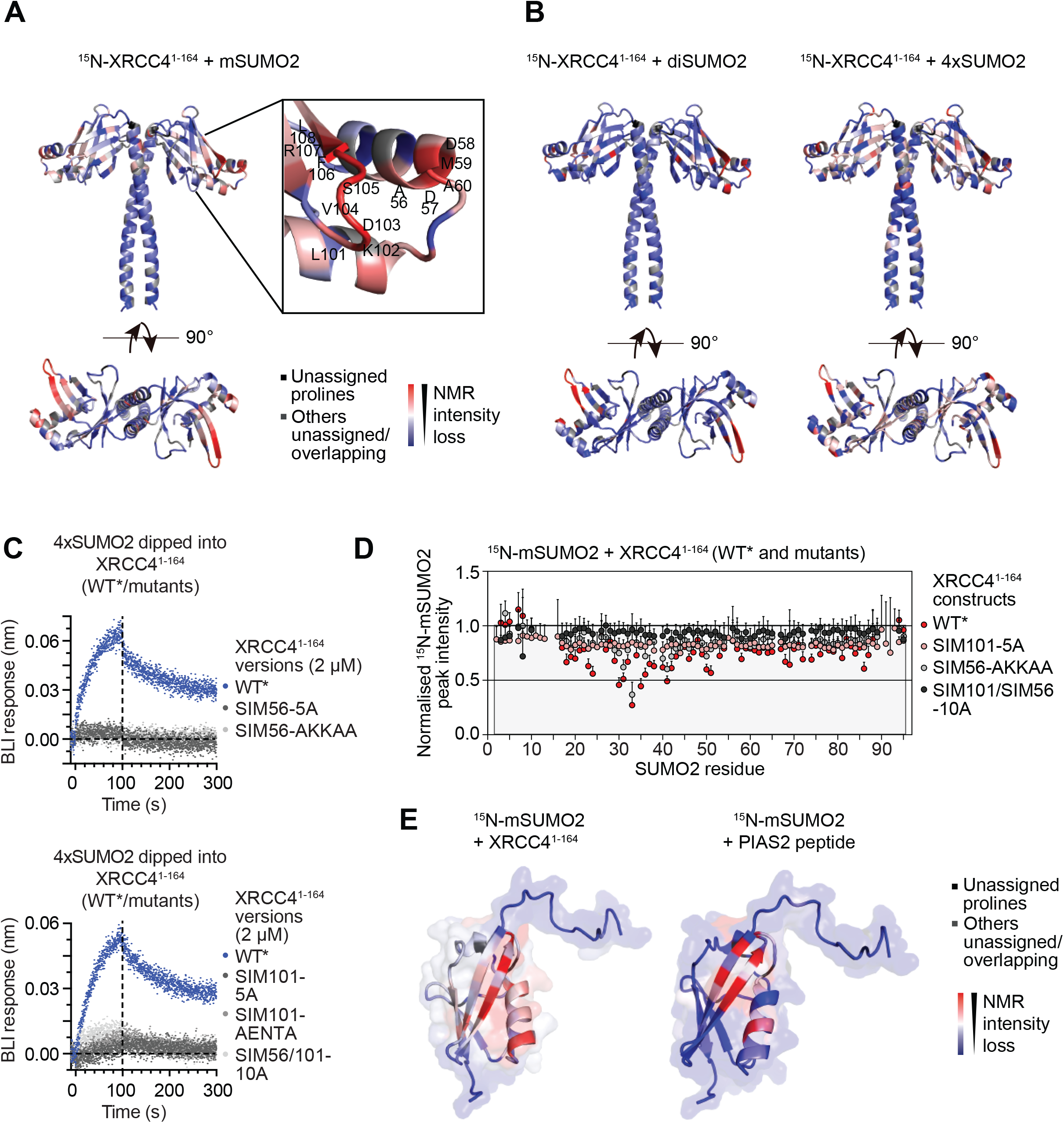
Unconventional SUMO2-specific binding of XRCC4 head domain. (**A**) XRCC4^1-164^ residues implicated in SUMO2 monomer (mSUMO2) binding, as indicated by intensity losses in the ^1^H-^15^N BEST-TROSY spectra of XRCC4^1-164^ after addition of increasing concentrations of mSUMO2. Colour gradient for the XRCC4 structure (PDB 1IK9) ranges from red (most affected by binding) to blue (unaffected by binding). For details see Methods section. Detailed view of key XRCC4^1-164^ residues affected is shown on the right. Unassigned prolines displayed in black, other unassigned/overlapping residues in grey. (**B**) XRCC4^1-164^ residues implicated in SUMO2 dimer (diSUMO2) (left) and SUMO2 tetramer (4xSUMO2; right) binding, as indicated by intensity losses in the ^1^H-^15^N BEST-TROSY spectra of XRCC4^1-164^ after addition of increasing concentrations of diSUMO2 or 4xSUMO2, respectively. Colour gradients for XRCC4 structures (PDB 1IK9) as indicated in (A). For details see Methods section. Unassigned prolines shown in black, other unassigned/overlapping residues in grey. (**C**) Mutating XRCC4 regions implicated in SUMO2 binding in XRCC4 head region, individually or combined, abrogates 4xSUMO2 binding to XRCC4^1-164^ as assessed by biolayer interferometry (BLI). Association and dissociation phases are separated by vertical dashed lines, as indicated. Mutations are explained at the end of the legend. (**D**) ^1^H-^15^N-TROSY peak ratios of mSUMO2 residues after addition of XRCC4^1-164^ mutants at 1:4 ratio to mSUMO2, compared to wildtype (WT) XRCC4^1-164^. Mutations are explained at the end of the legend. Error bars represent 1 standard deviation from the plotted value, as calculated from the noise levels in the TROSY spectra using the standard error propagation formula. Mutations are explained at the end of the legend **(E**) mSUMO2 residues implicated in XRCC4^1-164^ (left) and PIAS2 (right) binding, as indicated by intensity losses of the ^1^H-^15^N HSQC spectra of mSUMO2 after addition of increasing concentrations of XRCC4^1-164^ or PIAS2 peptide (467-VDVIDLTIESS-478). Colour gradients for mSUMO2 structures (PDB 2N1W) range from red (most affected by binding) to blue (unaffected by binding). For details see Methods section. Unassigned prolines shown in black, other unassigned/overlapping residues in grey. XRCC4^1-164^ mutations: SIM101-5A and SIM101-AENTA: XRCC4 residues 101-106 (LKDVS) mutated to alanines or AENTA, respectively; SIM56-5A and SIM56-AKKAA: XRCC4 residues 56-61 (ADDMA) mutated to alanines or AKKAA, respectively; SIM56/101-10A: XRCC4 residues 56-61 (ADDMA) and 101-106 (LKDVS) mutated to alanines.

The surface implicated in SUMO2 binding forms a negatively charged surface with a positively charged patch on one side (Supplementary Figure S4E), differentiating this surface from that of established SIM classes^2,13^. Consistent with our NMR analyses, individual or combined mutation of SIM56 and SIM101 abrogated binding of XRCC4^1-164^ to 4xSUMO2 in BLI assays (Fig. 5C, Supplementary Fig. S5A) and NMR titrations (Fig. 5D Supplementary Fig. S5A, B), while not affecting the overall structural integrity of the mutated proteins (Supplementary Fig. S5C). We next investigated if SIM56 and SIM101 were not only essential but also sufficient for SUMO2 binding. To this end, we performed NMR titrations with a peptide covering SIM101, the predominant XRCC4 SIM out of the two SIMs we identified. The lack of detectable binding, even at high molar equivalents of the peptide, illustrated that SIM101 was not sufficient for SUMO binding (Supplementary Fig. S5D; for positive peptide binding, see PIAS2 experiments below). These findings are consistent with neither wildtype nor mutant SIM101 peptide being able to compete with SUMO2 binding to XRCC4 in GFP-XRCC4 pulldown assays (Supplementary Fig. S5E). SIM56 was less affected than SIM101 in our NMR titrations (Supplementary Fig. S4A), and its mutation caused milder disruption of SUMO2 binding compared to SIM101 mutation, making it unlikely to function as an independent SIM (Fig. 5D; Supplementary Fig. S5B). We conclude that SIM56 and SIM101 likely synergise to facilitate SUMO2 binding of XRCC4, in agreement with their close spatial proximity (Fig. 5A). Collectively, these results uncover a non-conventional paralogue-specific SUMO2-binding mode mediated by the head domain of XRCC4.

Conventional SIMs can bind to different surfaces on SUMO1 and SUMO2^2,16^. To analyse how XRCC4-targeted SUMO2 surfaces correlate to the ones targeted by other SUMO receptors, we compared the ^1^H-^15^N HSQC spectra of mSUMO2 in the absence and presence of increasing concentrations of XRCC41-164. The analyses revealed binding of the XRCC4 head to the β_2_-strand and α_1_-helix of SUMO2, overlapping with the groove targeted by other known SUMO receptors such as PIAS2^19,108,16^, albeit with differences in the exact residue involvement (Fig. 5E; Supplementary Fig. S6A, B). Notably, several key residues of SUMO2, affected by XRCC4 binding, are not conserved in SUMO1, leading to alterations in charge (e.g. R36 in SUMO2 versus M40 in SUMO1) as well as in size (e.g. V23 in SUMO2 versus I27 in SUMO1; Supplementary Fig. S6C). These differences are reflected in a change in charge distribution across the corresponding SUMO2 and SUMO1 surfaces. For example, the positive charges on the XRCC4-bound SUMO2 surface are relatively weak and evenly distributed (Supplementary Fig. S6D), matching the homogenously distributed negative charges on the reciprocal XRCC4 surface (Supplementary Fig. S4E). By contrast, the equivalent SUMO1 surface possesses a strongly positively charged patch on one side, likely unfavouring binding of SUMO1 to XRCC4. Collectively, these analyses suggested that SUMO2 binding to XRCC4 may be stabilised by ionic interactions to help achieve paralogue specificity.

### SUMO2 interaction of XRCC4’s head domain is incompatible with XLF binding

Because the SUMO2 interaction surface on the head domain of XRCC4 overlaps with XRCC4 binding to another NHEJ core factor, XLF (Fig. 6A), we next investigated if SUMO2 and XLF binding were compatible with each other. To this end, we performed docking simulations using HADDOCK combined with 5 ns worth of molecular dynamics using GROMACS (2 fs steps). Two models indicative of SUMO2 binding to XRCC4 in opposing directions were retrieved that were consistent with the NMR intensity losses we used to generate the interaction restraints for HADDOCK (Fig. 6B; Supplementary Fig. S7A). Several surface-exposed hydrophobic residues (L101, V104 and F106) of SIM101 located in the β_6_-β_7_ hairpin of XRCC4, specifically in β_7_ and the short loop connecting β_6_ with β_7_, were key to both models, stacking up with a hydrophobic patch centred on F32 on the reciprocal SUMO2 surface (Fig. 6B). As expected, further stabilisation was mediated by ionic interactions e.g. between D103 (XRCC4) and K42 (SUMO2) in model 1, and between D57 (XRCC4) and K42 (SUMO2) in model 2 (Fig. 6B). Potential binding of SUMO2 in opposing directions raised the possibility that polySUMO2 chains could span the XRCC4 head domain with two of the SUMO moieties of individual chains binding to different XRCC4 monomers on each side of XRCC4’s dimeric head domain. Based on these simulations a minimum chain length of four SUMO2 moieties was required to facilitate the interaction of an N-terminal SUMO2 with XRCC4 on one side (via model 2), and a C-terminal SUMO2 on the other (via model 1; Supplementary Fig. S7B). Due to the marked spatial overlap between SUMO2, and XLF binding, to XRCC4, we predicted that binding of these two factors to XRCC4 would be mutually exclusive (Fig. 6C). In agreement with this hypothesis, mutation of SIM56 and SIM101, which abrogated XRCC4 binding to SUMO2 (see above) also abolished XRCC4 binding to XLF (Fig. 6D; Supplementary Fig. S7C). Indeed, XLF binding in these mutants was reduced to a level comparable to that of a 2KE XRCC4 mutant (K65 and K99 mutated to glutamic acids), a known XLF binding-deficient mutant (Fig. 6D; Supplementary Fig. S7C)^109,92^. Strikingly, the 2KE mutation rendered XRCC4^1-164^ also incapable of interacting with 4xSUMO2 (Fig. 6E), without affecting the overall structural integrity of the mutant (Supplementary Fig. S7D). Moreover, 4xSUMO2 showed a potential trend of competing with the XRCC4-XLF interaction when added to whole cell extracts at a final concentration of 1 μM (=25 μg) before performing GFP-XRCC4 pulldowns (~a quarter of reduction; Fig. 6F). Altogether, these findings supported a SUMO2-binding model that is incompatible with simultaneous binding of XLF to XRCC4.

**Figure 6.**
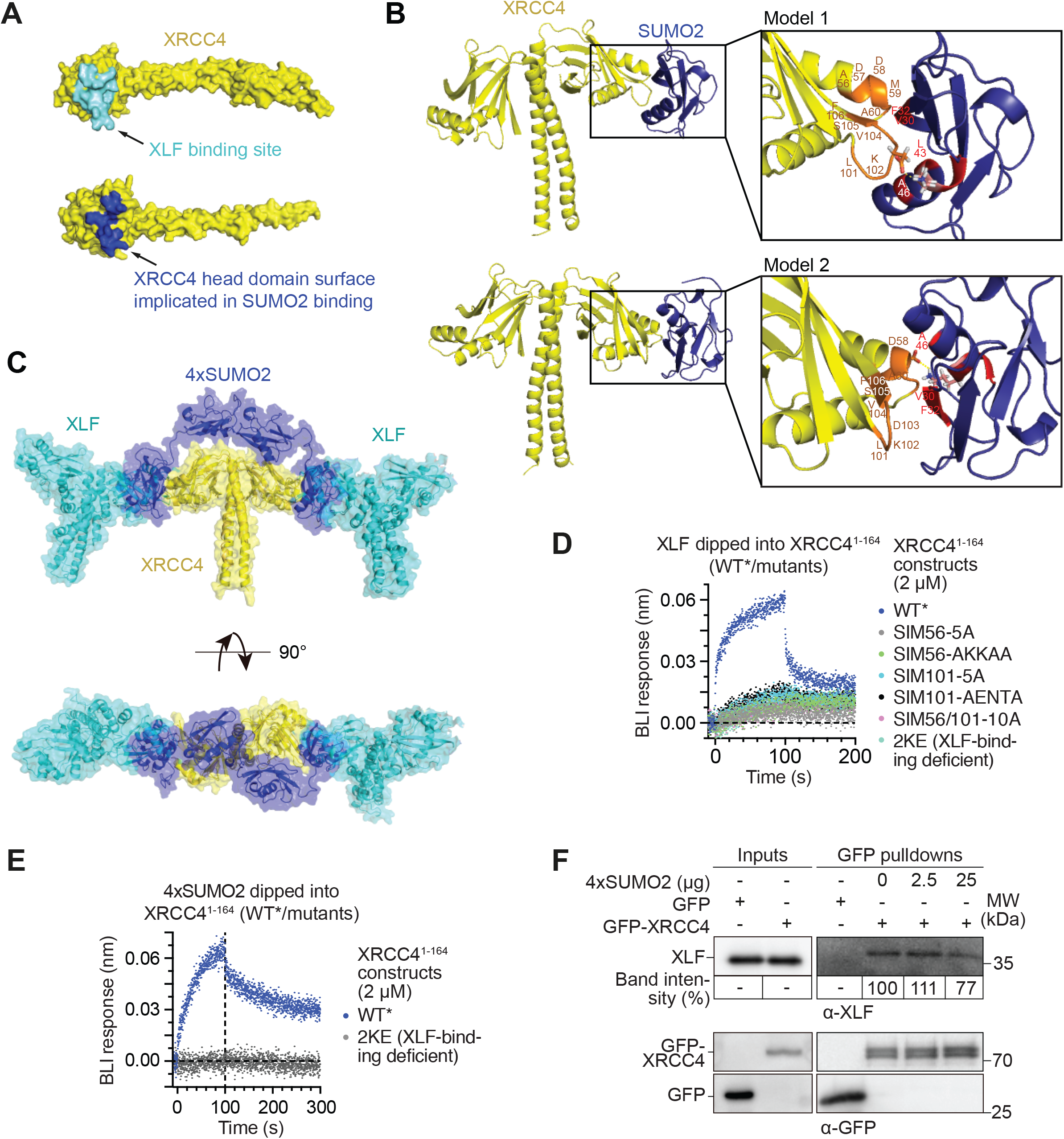
XRCC4 interaction with SUMO2 is incompatible with XLF binding. (**A**) XLF and SUMO2 binding to overlapping regions on XRCC4 (PDB 1FU1)^77^. XLF-binding region (top) encompasses XRCC4 residues within 5 Å distance of L115 of XLF, based on the crystal structure of the XRCC4-XLF complex (PDB 3RWR)^90^. SUMO2-binding region depicted on XRCC4 entails residues 56-60 and 101-108, according to the most strongly affected residues shown in Figure 5A and Supplementary Figure S4A. (**B**) Models of 4xSUMO2 (PDB 2D07)^133^ interaction with XRCC4 (PDB 1FU1) head domain, generated using HADDOCK and equilibrated with molecular dynamics using GROMACS. 56-ADDMA-60 and 101-LKDVSF-106 on XRCC4, and 30-VQFK-33, 40-LSKL-43 and A46 on SUMO2 highlighted in orange and red/pink, respectively, in models 1 and 2; polar interactions between D103 (XRCC4) and K42 (SUMO2) side chains in model 1, and D57 (XRCC4) and K42 (SUMO2) in model 2 highlighted with yellow dashed line. (**C**) XRCC4-XLF complex (PDB 3RWR)^90^ overlaid with XRCC4-SUMO2 interaction model according to Supplementary Figure S7B. (**D**) Biolayer interferometry (BLI) assays, illustrating abrogated XLF binding of the following SUMO2-binding deficient XRCC4^1-164^ mutants: SIM101-5A and SIM101-AENTA (XRCC4 residues 101-106 (LKDVS) mutated to alanines or AENTA, respectively); SIM56-5A and SIM56-AKKAA (XRCC4 residues 56-61 (ADDMA) mutated to alanines or AKKAA, respectively); and SIM56/101-10A (XRCC4 residues 56-61 (ADDMA) and 101-106 (LKDVS) mutated to alanines). Note that a combination of two XRCC4 mutations known to abrogate binding of full-length XRCC4 to XLF (2KE; K to E mutations at K65 and K99)^109,92^ also rendered XRCC4^1-164^ incapable of binding to XLF, and served as a positive control. Association and dissociation phases are separated by a vertical dashed line, as indicated. (**E**) 4xSUMO2 binding of the 2KE XLF binding-deficient XRCC4^1-164^ mutant described in (D) is abrogated in BLI assays. Association and dissociation phases are separated by a vertical dashed line, as indicated; BLI data for WT XRCC4^1-164^ are replicated from Figure 5C, forming part of the same experiment. (**F**) Precipitation of XLF by GFP-Trap pulldowns of GFP-XRCC4, ectopically expressed in HEK293T cells, from whole cell extracts after addition of increasing amounts of 4xSUMO2. Band intensities are averaged from two independent biological replicates and normalised to pulled-down XRCC4 levels.

### SUMO2 binding to XRCC4 coiled-coil is incompatible with LIG4 interaction

In addition to SUMO2 binding to the XRCC4 head domain, our carbene footprinting and XRCC4 truncation studies pointed towards a SUMO2-binding region in the XRCC4 coiled-coil located in or around the 170-178 region. Having successfully assigned the majority of residues in XRCC4^1-164^, we extended our NMR analyses to XRCC4^1-180^. Taking the XRCC4^1-164^ assignment as a basis we were able to assign the majority of the additional 16 residues and detected intensity loss and/or perturbation shifts after addition of mSUMO2 in some, but not all, of the residues, including E163, S167 and A168 (Fig. 7A, B). K164, C165 and V166 could not be assigned. However, given their sequence positioning in the centre of the most significantly affected residues (Fig. 7A), combined with the known importance of valines for mediating SUMO-SIM interactions^13,102^, we speculate that these residues contribute to SUMO2 binding.

**Figure 7.**
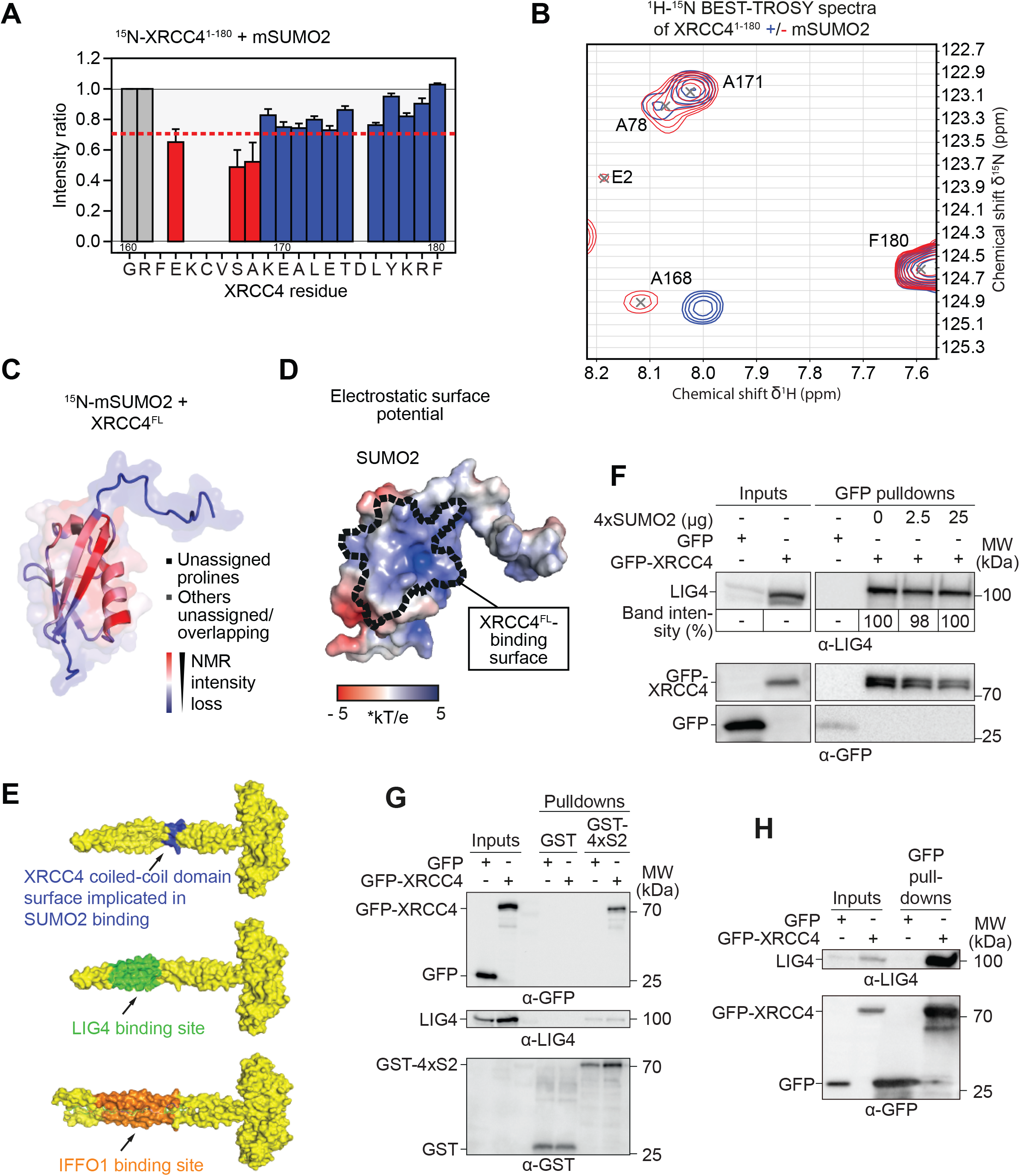
Non-conventional SUMO2-binding to XRCC4 coiled-coil overlaps with LIG4 and IFFO1 binding. (**A**) XRCC4^1-180^ residues implicated in SUMO2 monomer (mSUMO2) binding, as indicated by intensity losses in the ^1^H-^15^N BEST-TROSY spectra of XRCC4^1-180^ after addition of increasing concentrations of mSUMO2. Error bars represent 1 standard deviation from the plotted value, as calculated from the noise levels in the TROSY spectra using the standard error propagation formula. Grey bars indicate residues assigned in XRCC4^1-180^ with too low intensity for reliable measurement (arbitrarily set to 1); lack of bars represents residues that could not be assigned. Red and blue bars indicate assigned residues in XRCC4^1-180^ with reliable intensity measurements and levels of intensity ratio loss of more or less than 30%, respectively. (**B**) Representative panel of ^1^H-^15^N BEST-TROSY spectra of XRCC4^1-180^ without (red) and with (blue) 1 molar equivalent of mSUMO2 added. (**C**) mSUMO2 residues implicated in full-length XRCC4 (XRCC^FL^) binding, as indicated by intensity losses in the ^1^H-^15^N HSQC spectra of mSUMO2 after addition of 0.33 molar equivalents of XRCC4^FL^. Colour gradient for mSUMO2 structure (PDB 2N1W) ranges from red (most affected by binding) to blue (unaffected by binding). For details see Methods section. Unassigned prolines shown in black, other unassigned/overlapping residues in grey. (**D**) Electrostatic surface potential of SUMO2 created using APBS electrostatics plugin for Pymol^134^. Dashed line highlights key residues implicated in binding to XRCC4^FL^ according to (B) and Supplementary Figure S8A. Electrostatic potential values are multiples of kT/e, where kb is the Boltzmann's constant, T the temperature the calculation was run at (300 K) and e the charge of an electron, leading to a conversion factor of 25.85 mV. (**E**) SUMO2 (top), LIG4 (middle), and IFFO1 (bottom) binding to overlapping or spatially proximal regions on XRCC4 coiled-coil (PDB 1IK9 for top and middle and PDB 6ABO^81^ for bottom). SUMO2-binding region depicted on XRCC4 entails 163-EKCVSA-168 comprising residues with >30% intensity loss as well as their unassigned neighbouring residues according to (A). LIG4- and IFFO1-binding regions encompass XRCC4 residues 173-195 and 162-196, respectively^78,81^. (**F**) Precipitation of LIG4 by GFP-Trap pulldowns of GFP-XRCC4, ectopically expressed in HEK293T cells, from whole cell extracts after addition of increasing amounts of 4xSUMO2. Band intensities are averaged from two independent biological replicates and normalised to pulled-down XRCC4 levels. (**G**) GFP-XRCC4 pulled down with GST-4xSUMO2 from HEK293T whole cell extracts contains negligible amounts of LIG4. Inputs were 4% of the total. (**H**) GFP-Trap pulldowns of GFP-XRCC4 expressed in HEK293T cells contain a substantial fraction of the total LIG4 pool available in cells. Inputs were 4% of the total. S2: SUMO2.

Interestingly, the interaction surface on SUMO2 did not markedly change based on intensity losses in the ^1^H-^15^N HSQC spectra of mSUMO2 after addition of full-length XRCC4 compared to XRCC4^1-164^, apart from highlighting a small number of additional residues adjacent to previously implicated ones (compare Fig. 7C and Supplementary Fig. S8A to Fig. 5E and Supplementary Fig. S6A). We conclude that SIM56/SIM101 and SUMO-interacting residues on the coiled-coil bind to comparable interaction surfaces on SUMO2 with similar surface charge profiles (compare Fig. 7D to Supplementary Fig. S6D).

Strikingly the affected coiled-coil region overlaps with XRCC4 binding to two other proteins important for NHEJ: LIG4 and IFFO1 (Fig. 7E). In contrast to XRCC4 binding to XLF, addition of 4xSUMO2 to whole cell extracts did not compete with LIG4 binding of XRCC4 in pulldown assays even at high concentrations (compare Fig. 7F with Fig. 6F). These findings are in line with XRCC4 forming a stable, high-affinity complex with LIG4, based on an extensive interaction surface^82^. In light of the overlap between the SUMO2- and LIG4-binding sites we next tested if binding of the two proteins to XRCC4 was likely to be compatible. To this end, we performed GST-4xSUMO2 pulldown assays, demonstrating that XRCC4, but not LIG4, could be co-precipitated from whole cell extracts (Fig. 7G), while LIG4 was successfully co-precipitated from whole cell extracts in GFP-XRCC4 pulldowns (Fig. 7H). Collectively, these data suggested that LIG4 and polySUMO2 binding to XRCC4 were incompatible, and that similar principles may also apply to polySUMO2 and IFFO1 binding.

## Discussion

Although SUMOylations affect thousands of proteins regulating a gamut of cellular processes, only several tens of SUMO receptors decoding these SUMOylations have been validated. Establishing a pipeline based on human proteome microarrays and fluorescently labelled SUMO topologies, we uncover >200 new potential binary polySUMO2/3 receptors and validate a substantial fraction of them, markedly extending the known human SUMO receptor pool. Given the involvement of the identified receptors in diverse cellular pathways with no SUMO receptors previously assigned to them, these results serve as a platform for breaking new ground in SUMO biology. Indeed, numerous further opportunities now await exploration by exploiting our SUMO receptor screen to uncover mechanisms underlying established as well as understudied areas of SUMO biology, including epigenetics, pre-mRNA splicing, transcriptional regulation, DNA repair, cytoskeleton organisation and protein synthesis in health and disease, as well as in homeostasis and under cellular stress.

Our screening pipeline is widely applicable to other ubiquitin/UBL family members and we demonstrate its utility for identifying paralogue- and topology-specific binders, as well as for uncovering unprecedented ubiquitin/UBL-binding modes. To this end, we provide a paradigm for narrowing down ubiquitin/UBL-binding regions for proteins lacking conventional binding modes, using a recently developed chemical biology technique – carbene footprinting – that can be applied to proteins-of-interest independently of their size, amount of starting material available, and the affinity of the interaction, factors commonly limiting other analytical methods geared towards this purpose^105–107^.

By combining carbene footprinting and mutational studies with high-resolution structural analyses, we characterise XRCC4 as the first core NHEJ factor with SUMO receptor functions, revealing two distinct non-conventional SUMO-binding modules along its sequence. NHEJ has long remained understudied for its regulation by SUMOylation compared to other DSB and DNA repair pathways^23,24,110^. Intriguingly, the identified SUMO-binding mode of XRCC4 featured paralogue selectivity for SUMO2/3 over SUMO1 and for polySUMO chains over shorter topologies. While SUMO-binding relied on a hydrophobic patch in line with conventional SIMs, a positive charge at the core of the SUMO-binding module, K102, represented an unprecedented characteristic for this type of binding. Given the importance of acidic residues for mediating interactions with SUMO1^16^, this positive charge along with additional characteristics likely contributed to the observed paralogue specificity. Another intriguing finding arising from our studies was that the two SUMO-binding modules resided in highly structured domains of XRCC4, including α-helices rather than the commonly disordered regions of SIMs that tend to form β-strands upon SUMO binding^16^. Moreover, SUMO binding on XRCC4’s head domain was facilitated by two motifs acting in synergy with each other, conceptually reminiscent of the split-SIM interaction between TDP2 and SUMOylated TOP2^111^, albeit with different underlying molecular and stereospecific features. These findings suggest that spatially coherent surfaces formed by regions distal in their primary amino acid sequence may act as SUMO engagement platforms more commonly than anticipated. Taking this further, it will be interesting to investigate if coherent SUMO-binding modules could also be formed by different proteins, in analogy to group SUMOylations of protein complexes acting as integrated docking platforms for downstream receptors as recently proposed^25^. Despite the surprising features of XRCC4’s SUMO-binding surface, the reciprocal region on SUMO2 was similar to the one targeted by other SUMO receptors, albeit with different nuances in the precise residue involvement. These data highlight the versatility of SUMO to utilise the same region for interactions with a wide range of receptors relying on distinct binding modes.

Notably, XRCC4-like SIM features are significantly enriched in known SUMO receptors and can be detected in a large number of proteins overall, raising the possibility that XRCC4-like SUMO binding may be conserved across diverse areas in cell biology. Together with validating a receptor lacking both conventional and XRCC4-like SIMs – TCEAL6 – our results suggest an unanticipated spectral plasticity of SUMO-binding modes that has remained undiscovered, and for which our screen of binary SUMO receptors provides a rich resource.

Given the location of the SUMO-binding regions on XRCC4’s head and coiled-coil domains, different models of how polySUMO2 chains could bind to XRCC4 can be envisioned, including chains spanning across XRCC4’s head domain, and/or connecting its head and coiled-coil regions. Mechanistically, both SUMO-binding regions overlap or involve common features with other XRCC4 interaction regions important for different aspects of NHEJ. In this context, we note that the use of common interaction sites has emerged as an intriguing concept to make efficient use of a limited number of binding sites available on NHEJ core factors, thereby providing functional redundancy and/or diversity that can help cells deal with different types of DNA damage arising in distinct chromatin contexts and with varying sets and/or levels of functional repair factors available^112^. Such scenarios could apply to, and be relevant for, different tissues and developmental stages, as well as in different cancer settings due to differential regulation of NHEJ factors, their downregulation and/or their dysfunctioning.

Linking common binding sites to recognition of SUMOylations occurring in a spatiotemporally regulated manner such as in response to DNA damage^24^, could help cells coordinate the use of common binding regions in an optimal manner, enabling them to target distinct repair complexes to the most appropriate types of DSBs in varying chromatin environments and at different repair stages, while preventing harmful competition between them. Taking the above into consideration, we speculate that SUMO binding of XRCC4 may act as a back-up or complementary pathway to XRCC4-XLF, XRCC4-IFFO1 and/or XRCC4-LIG4-mediated functions depending on cellular context and without directly competing with them. The latter is consistent with the only modest or absent ability of SUMO to compete with XLF and LIG4 binding to XRCC4. In this regard, we note that depending on cellular context, various XLF redundancies have been described for example with PAXX, CYREN (also known as MRI), ATM, H2AX, MDC1 and 53BP1^113–117^. Our findings provide possible future avenues for exploring the mechanistic basis of such redundancies, which remain a puzzling phenomenon in NHEJ. Any such mechanisms are likely independent of XLF, IFFO1 and/or LIG4 due to their predicted incompatibility with SUMO to simultaneously bind to XRCC4.

Another possibility is that disruption of XRCC4 complexes by SUMO binding after completion of repair could be important for finalizing NHEJ, in analogy to the release of Ku after repair has taken place^118–121^. In that way, XRCC4-SUMO binding may contribute to other mechanisms negatively regulating XRCC4 interactions^101,122,123^. Additional SUMO-independent contact points between XRCC4 and its upstream SUMOylated protein(s), leading to an increased binding affinity, would make such a scenario more likely. Altogether, our results suggest further mechanistic studies to elucidate the precise events whereby XRCC4-SUMO binding regulates NHEJ processes. To this end, it will be interesting to identify and characterise the SUMO-dependent interactions of XRCC4 with its upstream SUMOylated protein(s).

Finally, we note that our work may have medical applications as targeting DDR and ubiquitin/UBL system components can be exploited to treat cancer^124,125^. Indeed, targeting NHEJ at the level of XRCC4 interactions represents an attractive and actively pursued approach to sensitise cancer cells, commonly displaying cryptic DNA repair pathway defects including NHEJ, via synthetic lethality and/or other mechanisms^126,127^. Similarly, given the importance of DSB repair pathway choice for determining CRISPR-Cas9 genome editing outcomes, targeting specific XRCC4 interactions important for NHEJ may also be relevant for increasing the efficiency of precise gene editing relying on homology-dependent repair^128^.

## Supplementary Figure legends

**Figure S1. Recombinant proteins used for polySUMO2 receptor validation.** Representative protein stains of the purified proteins used for Figure 2B.

**Figure S2. SUMO/ubiquitin topologies and interaction of XRCC4 with SUMO2 fused to a model substrate in human cells.** (**A**) Representative protein stains and immunoblots of SUMO topologies used for Figure 3C. Di-/polySUMO2 chains represent enzymatically linked chains via SUMO2’s internal K11 residue. The 4xSUMO2 is a genetically encoded linear fusion product of an N-terminal full-length SUMO2 linked to truncated SUMO2s (residues 11-92) for the 2^nd^, 3^rd^ and 4^th^ SUMO, and fused to an N-terminal 6xHis and a C-terminal Strep tag. (**B**) Representative protein stain of polySUMO3 chains used for Figure 3D (chains enzymatically linked via SUMO3’s internal K11 residue). (**C**) Representative protein stains of ubiquitin topologies used for Figure 3E, with tetra-ubiquitin chains enzymatically linked via ubiquitin’s internal lysines K6, K11, K29, K33, K48 and K63, or via its N-terminal methionine (M1). (**D**) XRCC4 interacts with SUMO2 fused to a model substrate (S; tet repressor) in human cells. Pulldowns of XRCC4 (GFP-XRCC4) and S-SUMO2 fusion proteins (SUMO2 N-terminally tagged with mCherry), ectopically expressed in HEK293T cells.

**Figure S3. XRCC4 binding to SUMO2 is independent of conventional SUMO interacting motifs (SIMs).** (**A**) Schematic of full-length XRCC4 (XRCC4^FL^), highlighting five putative SIMs (pSIMs) predicted in silico using JASSA and GPS-SUMO^13,73^. Underscores in the pSIM sequences in the bottom panel indicate residues forming extensive interactions with nearby XRCC4 residues, or residues buried deep inside the head domain of the XRCC4 dimer, as indicated in the XRCC4^1-164^ structure on the right (PDB 1IK9; colour code according to pSIMs on the left). (**B**) Consensus motifs for conventional SIMs according to JASSA^13^, created using PSSMSearch^135^. (**C**) Equilibrium analysis of SPR response unit maxima of polySUMO2 (top) and polySUMO3 (bottom) binding to immobilised XRCC4^FL^ wildtype (WT) and pSIM-5A mutants, which have the 5 amino acids of the pSIMs, as indicated in (B), mutated to alanines. (**D**) Protein stain and anti (α)-XRCC4 immunoblot of XRCC4^FL^ WT and pSIM-5A mutants as described in (C). (**E**) Equilibrium analysis of SPR response unit maxima of XLF binding to immobilised wildtype (WT) and 5A mutant XRCC4^FL^. XLF injected over immobilised WT and pSIM-5A mutants as described in (C). (**F**) Protein stain of purified recombinant XLF used for (E). (**G**) Equilibrium analysis of SPR response unit maxima of polySUMO2 binding to immobilised XRCC4^FL^ and XRCC4^1-213^. For representative protein stains of XRCC4^FL^ and XRCC41-213, see Figure 4D. K_D_: dissociation constant; RU: response unit. SPR: surface plasmon resonance.

**Figure S4. XRCC4^1-164^ residues implicated in binding to different SUMO topologies and electrostatic surface potential of XRCC4^1-164^-binding region to SUMO2.** (**A**) Top: XRCC4^1-164^ residues implicated in SUMO2 monomer (mSUMO2) binding, as indicated by intensity losses in the ^1^H-^15^N BEST-TROSY spectra of XRCC4^1-164^ after addition of increasing concentrations of mSUMO2. Colour gradient for XRCC4 residues ranges from red (most affected by binding) to blue (unaffected by binding). For details see Methods section. Unassigned prolines shown in black, other unassigned/overlapping residues in grey. Bottom: representative panels of XRCC4 ^1^H-^15^N BEST-TROSY spectra with key residues affected displayed in the bottom without (red) and with (blue) 0.5 monomer equivalents of mSUMO2 added. (**B**) XRCC4^1-164^ residues implicated in SUMO2 dimer (diSUMO2) binding, as indicated by intensity losses in the ^1^H-^15^N BEST-TROSY spectra of XRCC4^1-164^ after addition of increasing concentrations of diSUMO2. Colour gradient for XRCC4 residues ranges from red (most affected by binding) to blue (unaffected by binding). For details see Methods section. Unassigned prolines shown in black, other unassigned/overlapping residues in grey. (**C**) XRCC4^1-164^ residues implicated in SUMO2 tetramer (4xSUMO2) binding, as indicated by intensity losses in the ^1^H-^15^N BEST-TROSY NMR spectra of XRCC4^1-164^ after addition of increasing concentrations of 4xSUMO2. Colour gradient for XRCC4 residues ranges from red (most affected by binding) to blue (unaffected by binding). For details see Methods section. Unassigned prolines shown in black, other unassigned/overlapping residues in grey. (**D**) XRCC4^1-164^ residues implicated in SUMO1 monomer (mSUMO1) binding, as indicated by intensity losses in the ^1^H-^15^N BEST-TROSY spectra of XRCC4^1-164^ after addition of increasing concentrations of mSUMO1. Colour gradient for XRCC4 residues ranges from red (most affected by binding) to blue (unaffected by binding). For details see Methods section. Unassigned prolines shown in black, other unassigned/overlapping residues in grey. (**E**) Electrostatic surface potential (left) of XRCC4 head domain (PDB 1IK9; ; orientation indicated on the right) with key residues involved in SUMO2 binding (56-ADDMA-60, 101-LKDVS-105) highlighted inside dashed line, created using APBS^134^. Positively charged surfaces are shown in blue, neutral ones in white, negatively charged in red with values displayed as multiples of kT/e, where kb is the Boltzmann's constant, T the temperature of the calculation (300 K) and e the charge of an electron, leading to a conversion factor of 25.85 mV.

**Figure S5. Protein stains, SUMO binding and folding status of SIM56/SIM101 XRCC4^1-164^ mutants, and SIM101 peptide analyses.** (**A**) Representative protein stains of recombinant proteins used for Figure 5. For recombinant 4xSUMO2 protein stain see Supplementary Figure S2. (**B**) ^1^H-^15^N HSQC spectra peak intensity ratios of mSUMO2 residues after addition of XRCC4^1-164^ mutants as indicated in 1:4 ratio to mSUMO2, compared to WT XRCC4^1-164^. SIM56-5A: 56-ADDMA-60 of XRCC4 head domain mutated to AAAAA; SIM101-AENTA: 101-LKDVS-105 of XRCC4 head domain mutated to AENTA. WT intensities replicated from Figure 5D as reference. Error bars represent 1 standard deviation as calculated from the average noise level in the HSQC spectra, using the standard error propagation formula. (**C**) 1D ^1^H NMR spectra of wildtype (WT) and mutant XRCC4^1-164^, demonstrating no major effects on the folding status of the introduced mutations. Mutations are as follows: SIM101-5A and SIM101-AENTA: 101-LKDVS-105 of XRCC4 head domain mutated to AAAAA or AENTA, respectively; SIM56-5A and SIM56-AKKAA: 56-ADDMA-60 of XRCC4 head domain mutated to AAAAA or AKKAA, respectively; SIM56/SIM101-10A: 101-LKDVS-105 and 56-ADDMA-60 in XRCC4 domain mutated to alanines. (**D**) SIM101 is not sufficient for SUMO2 binding. Peak heights of ^15^N-labeled monomeric SUMO2 (mSUMO2) in ^1^H-^15^N HSQC spectra after addition of 1 equivalent of SIM101 peptide (LKDVSFRLGSF; XRCC4 residues 101-111) compared to WT XRCC4^1-164^ spectrum (replicated from Figure 5D) used as reference. (**E**) Pulldowns of GFP-XRCC4 and SUMO2 fusion proteins, ectopically expressed in HEK293T cells, in the absence or presence of SIM101 wildtype (WT; 99-KNLKDVSFRLGSF-111) and mutated (99-KNAAAAAFRLGSF-111) peptide. 200 μg of WT or mutated peptide were added to the cellular extracts prior to the incubation with the beads. SUMO2 was coupled to a model substrate (S; tet repressor) and fused to mCherry (S-mCherrySUMO2), with S-mCherry serving as control. Inputs were 4% of the total.

**Figure S6. XRCC4-binding region on SUMO2 and comparative SUMO paralogue features.** (**A**) Top: Intensity losses in ^1^H-^15^N HSQC spectra of monomeric SUMO2 (mSUMO2) after addition of increasing concentrations of XRCC4^1-164^. Colour gradient for XRCC4 residues ranges from red (most affected by binding) to blue (unaffected by binding). For details see Methods section. Unassigned prolines shown in black, other unassigned/overlapping residues in grey. Bottom: representative panel of mSUMO2 ^1^H-^15^N HSQC spectra displaying key residues affected without (red) and with (blue) 0.25 equivalents of XRCC4^1-164^ added. (**B**) ^1^H-^15^N HSQC spectra like described in (A) but for PIAS2 11-mer peptide (467-VDVIDLTIESS-478). (**C**) Sequence alignment of mature forms of human SUMO1, SUMO2 and SUMO3 paralogues according to Clustal Omega^136^ with secondary structure elements depicted below, based on PDB entries 2N1V^137^ for SUMO1, 2N1W for SUMO2, and 1U4A^138^ for SUMO3. (**D**) Electrostatic surface potentials of SUMO2 and SUMO1. Left: dashed lines highlight key residues of SUMO2 implicated in binding to XRCC4^1-164^ according to residues boxed in SUMO2 in (C). Right: equivalent SUMO1 residues as boxed in (C). Electrostatic potential values were plotted onto the surface using APBS^134^ with negatively and positively charged values ranging from red to blue, respectively. Values displayed are multiples of kT/e, where kb is the Boltzmann's constant, T the temperature the calculation was run at (300 K) and e the charge of an electron, leading to a conversion factor of 25.85 mV.

**Figure S7. Model of 4xSUMO2 binding to XRCC4 head domain and folding status of XRCC4^1-164^ 2KE mutant.** (**A**) Model of 4xSUMO2-XRCC4 interactions, highlighting the interaction restraints used for HADDOCK based on NMR intensity losses as follows: for XRCC4 (PDB 1IK9) “active” residues were 57, 59, 102, 103, 104, 105 and 106, “passive” residues were 56, 62, 65, 99, 101, 109 and 111; for SUMO2 (PDB 2D07) “active” residues were 30, 31, 33, 35, 40, 41, 42, 50 and 51, “passive” residues were 29, 36, 38, 43, 68, 84, 86. Model 1 is shown as an example with active residues highlighted in red, passive ones in orange. Restraints were similarly adhered to in model 2. (**B**) Model of 4xSUMO2 binding to XRCC4 head domain based on docking simulations using HADDOCK and equilibrated by molecular dynamics using GROMACS; docked protein structures based on PDB entries 1IK9 for XRCC4 (chain A) and 2D07 for SUMO2. (**C**) Representative protein stains of purified XLF-6xHis and XRCC4^1-164^ 2KE used for Figure 6D. For representative protein stains of wildtype and SIM56/SIM101-mutant XRCC4^1-164^ see Supplementary Figure S5A. (**D**) 1D ^1^H NMR spectra of wildtype (WT) XRCC4^1-164^ and XLF binding-deficient XRCC4 (2KE mutant with K65 and K99 mutated to glutamic acids)^109,92^, highlighting no marked changes in folding state of XRCC4 2KE compared to WT. XRCC4 WT spectrum replicated from Supplementary Figure S5C as reference.

**Figure S8. SUMO2 residues implicated in binding to full-length XRCC4.** (**A**) Intensity losses in ^1^H-^15^N HSQC spectra of mSUMO2 after addition of increasing concentrations of full-length XRCC4 (XRCC4^FL^). Colour gradient for XRCC4 residues ranges from red (most affected by binding) to blue (unaffected by binding). For details see Methods section. Unassigned prolines shown in black, other unassigned/overlapping residues in grey.

## Methods

### PolySUMO2 microarray staining and analysis

polySUMO2 chains (#ULC-220, Boston Biochem) were directly labeled with Cy3 following the manufacturer’s guidelines (GE Healthcare). After 45 min incubation in the dark, 10% reaction volume of 2 M Tris/HCl pH 7.5 was added to squelch unreacted dye, and the incubation extended in the dark for 10 min. polySUMO2 chains where then purified in PD25 spin columns according to the manufacturer’s recommendation (GE Healthcare). The purified and labeled polySUMO2 chains were immediately applied to blocked human proteome microarrays (HuProt™v2.0, CDI Laboratories). Microarrays were removed from 20°C storage and placed at room temperature (RT) for 15 min before opening, to avoid condensation. Arrays were then blocked for 1 h at RT in PBS containing 0.05% Tween-20, 20 mM reduced glutathione, 1 mM DTT, 3% BSA, and 25% glycerol. Three PBS washes preceded a 90-min incubation step at RT with labeled polySUMO2 chains (or Tris-squelched Cy3 dye as a reference). After two washing steps with PBS containing 0.05% Tween-20, two PBS washes, and two washes with water, centrifugal drying (1,000 rpm for 5 min at RT) was performed and the arrays scanned using a GenePix scanner (4100A by Molecular Devices). Microarray images were gridded and quantitated using GenePix Pro (v7) software. Median intensities (features and local backgrounds) were utilised, and signal-to-noise ratios calculated. Values were then normalised to biological controls within each array and duplicate features (representing identical proteins) summarised by average. These values were compared between arrays (polySUMO2 hybridised minus mock-treated array) then Loess transformed by print tip and location to remove technical sources of error^139^, resulting in the final estimate of magnitude change (M-value). T-test (paired, 2-tailed) was used to assess the statistical significance (*p*-value) of each estimate (under the null hypothesis that M=0). The threshold for proteins classifying as polySUMO2 receptor candidates was set to 1 standard deviation of the population above the population average. Given that the M-value is a twice-normalised (biologically and for technical sources of error) difference between mean signal-to-noise ratios generated from relative fluorescence units, it is reported/graphed as ‘M-value’ without units.

### Carbene footprinting

Samples were prepared and analysed as previously described^105^. Briefly, 20 μM full-length XRCC4 or 25 μM XRCC4^1-164^ were mixed with 20 μM or 25 μM of mSUMO2, respectively, in a buffer containing 20 mM HEPES pH 6.8, 140 mM NaCl, 1 mM EDTA, 2 mM DTT and 0.02% NaN_3_, as well as 10 mM of aryldiazirine probe (total volume, 20 μL). The mixture was left to equilibrate for 5 min at RT before 6 μL aliquots were placed in crystal-clear vials (Fisher Scientific UK) and snap-frozen in liquid nitrogen. The labelling reaction was initiated by photolysis of the mixture using the third harmonic of a Nd:YLF laser (Spectra Physics, repetition frequency 1,000 Hz, pulse energy 125 μJ) at a wavelength of 347 nm. The frozen samples were irradiated for 10 s. All experiments were performed in triplicate. Following irradiation, samples were thawed, reduced (10 mM DTT in 10 mM ammonium bicarbonate), alkylated (55 mM iodoacetamide in 10 mM ammonium bicarbonate) and incubated at 37°C with trypsin overnight (1:20 protease/protein ratio in 10 mM ammonium bicarbonate). The analysis of the digests was carried out on a Bruker MaXis II ESI-Q-TOF-MS connected to a Dionex 3000 RS UHPLC fitted with an ACE C18 RP column (100 x 2.1 mm, 5 μm, 30°C). The column was eluted with a linear gradient of 5–100% MeCN containing 0.1% formic acid over 40 min. The mass spectrometer was operated in positive ion mode with a scan range of 200– 3,000 m/z. Source conditions were: end plate offset at −500 V; capillary at −4,500 V; nebulizer gas (N_2_) at 1.6 bar; dry gas (N_2_) at 8 L/min; dry temperature at 180°C. Ion transfer conditions were: ion funnel RF at 200 Vpp; multiple RF at 200 Vpp; quadrupole low mass at 55 m/z; collision energy at 5.0 eV; collision RF at 600 Vpp; ion cooler RF at 50–350 Vpp; transfer time at 121 s; pre-pulse storage time at 1 μs. A previously described method was used to quantitate the fraction of each peptide modified^106^. Briefly, the chromatograms for each singly-labelled and unlabelled peptide were extracted within a range of ±0.1 m/z and the spectrum for each peak was manually inspected to ensure the sampling of the correct ion only. The peptide fractional modification was calculated using Equation X.

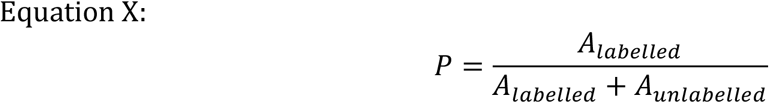

Where A_labelled_ and A_unlabelled_ correspond respectively to the peak area of each labelled and unlabelled peptide. Differences in the extent of labelling between peptides were considered significant when the *p*-value obtained from a Student t-test was <0.05.

### NMR spectra acquisition and analysis

Protein spectra were recorded at 310 K on a Bruker 800 MHz spectrometer with a ^1^H/^13^C-^15^N TCI cryoprobe equipped with z-gradients in 20 mM Hepes pH 6.8, 140 mM NaCl, 1 mM EDTA, 2 mM DTT, 100 mM arginine, 100 mM glutamic acid, 0.02% NaN_3_, unless otherwise specified. XRCC4 ^1^H-^15^N spectra were standard Bruker BEST-TROSY with phase-sensitive Echo/Antiecho gradient selection. SUMO ^1^H-^15^N spectra were standard Bruker sensitivity-enhanced, phase-sensitive HSQC spectra using Echo/Antiecho gradient selection. 1D ^1^H spectra were recorded using excitation sculpting water suppression. The assignments for mSUMO2 were taken from BMRB entry 6801 and temperature and buffer conditions were incremented from the conditions used in the assignment to those used for this study, to allow for the associated chemical shift changes. Assignment of 6xHis-XRCC4^1-164^ was done by the standard triple resonance methods^140^, using a ^2^H-^15^N-^13^C-labelled protein sample, with the amides back-exchanged to ^1^H. Back-exchange was spontaneous over approximately one week. 3D spectra were recorded using non-uniform sampling with a multidimensional Poisson Gap scheduling strategy with sinebell weighting^141^. The chemical shifts from the assignment were used to calculate secondary structure propensity using TALOS+^142^, which agreed with available crystal structures. 169 of the 189 backbone amides in the construct were assigned, with 15 of the 21 unassigned residues being in the 6xHis tag. Assignment of 6xHis-XRCC4^1-180^ was attempted by the same methods using a ^1^H-^15^N-^13^C-labelled protein sample. The majority of ^1^H^15^N crosspeaks were extremely low intensity, but in the same positions as in the 6xHis-XRCC4^1-164^ construct. Residues 167-180 were assigned. Assignments are deposited with BMRB code (to be determined). Data were processed and visualised using Topspin 3.5 (Bruker), and protein backbone assignment was done using CCPN Analysis 2.1^143^. Protein-protein interactions were analysed using CCPNAssign (v3.1)^144^.

### Colour gradients for NMR analyses

#### ^15^N-labelled XRCC4^1-164^ + mSUMO2

Colour gradient range for the XRCC4 structure (PDB 1IK9) and sequence based on intensity losses in the ^1^H-^15^N BEST-TROSY spectra of XRCC4^1-164^ after addition of increasing concentrations of mSUMO2 as follows: from red (most affected by binding: <45% intensity after addition of 0.1 monomer equivalents) to blue (unaffected by 4-fold excess over monomer), with intermediate points as follows: 45-66% intensity at 0.1 equivalents, <45% intensity at 0.5 equivalents, 45-66% intensity at 0.5 equivalents, <45% intensity at 1 equivalent (white), 45-66% intensity at 1 equivalent, <45% intensity at 4 equivalents and 45-66% intensity at 4 equivalents.

#### ^15^N-labelled XRCC4^1-164^ + diSUMO2

Colour gradient range for XRCC4 structure (PDB 1IK9) and sequence based on intensity losses in the ^1^H-^15^N BEST-TROSY spectra of XRCC4^1-164^ after addition of increasing concentrations of diSUMO2 as follows: from red (most affected by binding: <20% intensity after addition of 0.125 monomer equivalents), to blue (unaffected by 0.25 equivalents), with intermediate points as follows: 20-30% intensity, 30-45% intensity, 45-66% intensity at 0.125 equivalents; <20% intensity (white), 20-30% intensity, 30-45% and 45-66% intensity at 0.25 equivalents.

#### ^15^N-labelled XRCC4^1-164^ + 4xSUMO2

Colour gradient range for XRCC4 structure (PDB 1IK9) and sequence based on intensity losses in the ^1^H-^15^N BEST-TROSY spectra of XRCC4^1-164^ after addition of increasing concentrations of 4xSUMO2 as follows: from red (most affected by binding: <20% intensity on addition of 0.1 monomer equivalents) to blue (unaffected by 0.67 equivalents), with intermediate points as follows: 20-30% intensity, 30-45% intensity and 45-66% intensity at 0.1 equivalents; <20% intensity (white), 20-30% intensity, 30-45% and 45-66% intensity at 0.67 equivalents.

#### ^15^N-labelled XRCC4^1-164^ + mSUMO1

Colour gradient range for XRCC4 structure (PDB 1IK9) and sequence according to chemical shift perturbations (CSP) in the ^1^H-^15^N-TROSY NMR spectra of XRCC4^1-164^ after addition of 1 monomer equivalent of mSUMO1 as follows: from red (most affected by binding: >5 standard deviations intensity on addition of 1 monomer equivalents), to blue (unaffected by 1 equivalent), with intermediate points as follows: 4, 3, 2 and 1 standard deviation(s).

#### ^15^N-labelled mSUMO2 + XRCC4^1-164^

Colour gradient range for mSUMO2 structure (PDB 2N1W) and sequence based on intensity losses in the ^1^H-^15^N HSQC spectra of mSUMO2 after addition of increasing concentrations of XRCC4^1-164^ as follows: from red (most affected by binding: <20% intensity after addition of 0.25 mSUMO2 equivalents) to blue (unaffected by 1 equivalent), with intermediate points as follows: 21-30% intensity, <31-40% intensity and 41-66% intensity at 0.25 equivalents; <20% intensity (white), 21-30% intensity, 31-40% intensity and 41-66% intensity at 0.5 equivalents.

#### ^15^N-labelled mSUMO2 + XRCC4^FL^

Colour gradient range for mSUMO2 structure (PDB 2N1W) and sequence based on intensity losses in the ^1^H-^15^N HSQC spectra of mSUMO2 after addition of 0.33 molar equivalents of XRCC4^FL^ as follows: from red (most affected by binding: <20% intensity after addition of 0.33 mSUMO2 equivalents) to blue (unaffected by 0.33 molar equivalents), with intermediate points as follows: 21-30% intensity at 0.33 equivalents, 31-40% intensity at 0.33 equivalents, 41-66% intensity at 0.33 equivalents.

#### ^15^N-labelled mSUMO2 + PIAS2 peptide

Colour gradient range for mSUMO2 structure (PDB 2N1W) and sequence based on intensity losses in the ^1^H-^15^N HSQC spectra of mSUMO2 after addition of 0.5 mSUMO2 equivalents of PIAS2 peptide (467-VDVIDLTIESS-478) as follows: from red (most affected by binding: <20% intensity after addition of 0.5 mSUMO2 equivalents of PIAS2 peptide) to blue (unaffected by 0.5 equivalents), with intermediate points as follows: 21-30% intensity, <31-40% intensity and 41-66% intensity.

### Docking simulations

Docking simulations were performed using HADDOCK^145^. NMR intensity losses were used to generate ambiguous interaction restraints. The “active” residues were 57, 59, 102, 103, 104, 105 and 106 for XRCC4 and 30, 31, 33, 35, 40, 41, 42, 50 and 51 for mSUMO2. The “passive” residues were 56, 62, 65, 99, 101, 109 and 111 for XRCC4 and 29, 36, 38, 43, 68, 84, 86 for mSUMO2. The docked protein structures were based on PDB entries 1IK9 for XRCC4 (chain A) and 2D07 for SUMO2. In order to assess the reproducibility, the HADDOCK docking was repeated with one active residue omitted from each binding partner for each possible pair. The 63 docked structures generated, clustered into 7 classes, and these 7 clusters showed only 2 orientations of mSUMO2 relative to XRCC4. These two orientations were used to generate a model of 4xSUMO2 binding to XRCC4, which was then equilibrated by molecular dynamics using GROMACS (5 ns in 2 fs steps using the AMBER99SB-ILDN forcefield and TIP3P water). Submitted to PDB (to be determined).

### Surface plasmon resonance (SPR)

Experiments were performed on a ProteOn XPR36 instrument (BioRad Laboratories) using a running buffer containing 100 mM NaCl, 10 mM Hepes pH 7.0 and 0.1% (v/v) Igepal. Recombinant XRCC4-6xHis was immobilised on a GLC chip (BioRad Laboratories) in the vertical orientation. Chip channels were activated using a mixture of 25 mM *N*-ethyl-*N*′-(3-dimethylaminopropyl) carbodiimide (EDC) and 15 mM sulfo-*N*-hydroxysuccinimide (sulfo-NHS). Proteins were immobilised in 10 mM sodium acetate buffer, pH 4.5 (BioRad Laboratories). Remaining cross-linking sites were blocked by injection of 150 μL of 1 M ethanolamine–HCl (pH 8.5). Different SUMO or ubiquitin topologies were run in horizontal orientation at concentrations specified in each panel. Surface regeneration was accomplished with a pulse of 50 mM NaOH at 100 μL/min. All experiments were performed at 25°C. K_D_ values were obtained via non-linear regression based on a single-site binding model (GraphPad Prism v6.0h).

### Biolayer interferometry (BLI)

BLI was performed using an Octet RED96 instrument (ForteBio). 50 μg of recombinant 6xHis-4xSUMO2-Strep was biotinylated using EZ-link NHS-PEG4-Biotin (Thermo Fisher) following the manufacturer’s protocol. Excess biotin was removed using Zeba desalting spin columns (Thermo Fisher). 1 μg of biotinylated 6xHis-4xSUMO2-Strep were immobilised on streptavidin (SA) biosensors (ForteBio) until an approximately 1,000 nm response was reached. The baseline was set by submerging 4xSUMO2-captured sensors in kinetics buffer (PBS + 0.02% Tween-20, 0.1% BSA, 0.05% NaN_3_) in a 96-well plate, integrating an orbital shake function. The binding curves were obtained by dipping the sensors in 96-well plates containing the analytes diluted in kinetics buffer, or kinetics buffer only as reference. Finally, the sensors were dipped in fresh kinetics buffer for the dissociation step. Sensors were regenerated using 100 mM glycine pH 2.5 prior to reuse. Unloaded SA biosensors were used as controls, and as a reference where unspecific binding was observed.

### Cell Culture

HEK293T cells were cultured in high-glucose Dulbecco’s Modified Eagle Medium (DMEM, Sigma), supplemented with 10% (v/v) fetal bovine serum (FBS, Sigma), 2 mM L-glutamine, 100 U/mL penicillin, and 100 μg/mL streptomycin (Gibco) at 37°C in a humidified atmosphere at 5% CO_2_. Transfections were carried out using Fugene6 (Promega) according to the manufacturer’s instructions. Ionizing radiation (IR) treatments were performed using a CellRad Faxitron instrument (Faxitron Bioptics, LLC).

### Construct design

The GST-diSUMO2 plasmid was generated by overlapping PCRs of SUMO2 lacking the final two residues fused to a truncated SUMO2 (corresponding to amino acids 11-95) with a di-glycine linker in between the two SUMO2 and inserted in the BamHI/EcoRI sites of the pGEX-2T vector. XRCC4^1-180^ (cysteines 93, 128 and 130 mutated to alanines) for NMR purposes was amplified from the pET-28a-XRCC4^1-164^ (C-to-A) vector with a reverse primer containing the sequence for residues 165-180 and inserted in the pET-28a vector linearised with NcoI/EcoRI, and validated by sequencing. XRCC4^1-164^ SIM101-5A (cysteines 128 and 130 mutated to alanines), SIM101-AENTA (cysteines 128 and 130 mutated to alanines), SIM56-5A (cysteines 93, 128 and 130 mutated to alanines), SIM56AKKAA (cysteines 93, 128 and 130 mutated to alanines), SIM56/101-10A (cysteines 128 and 130 mutated to alanines) were generated by overlapping PCRs with mutagenic primers using the pET-28a-XRCC4^1-164^ as template . The XRCC4^1-164^ 2KE mutant was amplified from the full-length XRCC4 K65E K99E mutant^109^, a kind gift from Murray Junop (Western University London, USA) and inserted into the pET-28a vector linearised with NcoI/EcoRI, and validated by sequencing. XLF was amplified from pGEX2TKP-XLF^91^ and subcloned into the pHAT5 vector. XRCC4^1-172^, XRCC^1-180^, XRCC4^1-213^ and XRCC4^1-270^ were amplified from pHAT5-XRCC4^FL^. XRCC4 pSIM8, pSIM33, pSIM123 and pSIM181 were generated by overlapping PCRs with mutagenic primers using pHAT5-XRCC4^FL^ as a template and inserted in the NcoI/BamHI sites of the pHAT5 expression vector. The following plasmids were used for mammalian transfections: pEGFP-C1-FLAG-XRCC4, pEGFP-C1 (Clontech) containing FLAG-XRCC4 (denoted GFP-XRCC4), has been described previously^146^. pEGFP-C1-FLAG was generated by removing XRCC4 from pEGFP-C1-FLAG-XRCC4 using BamHI. pCAGGS-tetR_NLS_mCherry was a kind gift from Edith Heard (EMBL, Heidelberg, Germany)^147^. pCAGGS-tetR_NLS_mCherry-4xSUMO2 was cloned by inserting the 4xSUMO2 sequence, amplified from the 6xHis-4xSUMO2-Strep expression vector, and inserted in the BsrgI/EcoRI sites of pCAGGS-tetR_NLS_mCherry.

### Streptavidin pulldowns

6xHis-4xSUMO2-Strep was biotinylated as described above. 30 μg of biotinylated protein were mixed with 60 μL of streptavidin agarose resin (Thermo Fisher) and rotated (end-over-end) for 30 min at RT. The 4xSUMO2-bound beads were then centrifuged and washed. 30 μg of 6xHis-XRCC4 diluted in 500 μL of PBS containing 1% Igepal were added to the 4xSUMO2-captured beads and rotated (end-over-end) for 1 h at RT. Subsequently, the beads were washed 5 times in PBS containing 1% Igepal, and resuspended in 2x SDS Laemmli buffer (120 mM Tris-HCl pH 6.8, 4% SDS, 20% glycerol, 0.02% bromophenol blue and 2.5% β-mercaptoethanol). Proteins were visualised with InstantBlue stain (Expedeon).

### GFP-immunoprecipitations

HEK293T cells transfected with the desired expression construct were washed with ice-cold PBS and scraped into lysis buffer (50 mM Tris/HCl pH 7.5, 150 mM NaCl, 10% glycerol, 2 mM MgCl_2_, 10 mM N-ethylmaleimide) with 1x Complete EDTA-free protease inhibitors (Roche) and 6 μl benzonase (Millipore) and rotated at RT for 15 min. Subsequently, the lysates were centrifuged at 16,000 g for 60 min and the supernatant bound to 25 μl of GFP-trap magnetic beads (Chromotek) for 1 h with end-over-end rotation at 4°C. Protein-bound beads were then washed 5 times with lysis buffer and resuspended in 2x SDS Laemmli buffer (120 mM Tris-HCl pH 6.8, 4% SDS, 20% glycerol, 0.02% bromophenol blue and 2.5% β-mercatoethanol). For competition experiments the indicated amount of recombinant 6xHis-4xSUMO2-Strep was added to the lysates prior to the incubation with the beads. 4% of input lysate was loaded unless stated otherwise.

### GST-immunoprecipitations

Pulldowns with cellular extracts were carried out as described for GFP-immunoprecipitations using 1-2 μg of GST-fused proteins bound to glutathione magnetic beads (Promega). For immunoprecipitations of recombinant proteins, 1-2 μg of GST-fused proteins bound to glutathione magnetic beads were resuspended in 600 ul of binding buffer (50 mM Tris/HCl pH 7.5, 150 mM NaCl, 10% glycerol, 2 mM MgCl_2_, 10 mM N-ethylmaleimide) containing 2 μg or equimolar concentrations of His-tagged recombinant proteins and incubated for 1 h with end-over-end rotation at 4°C. Protein-bound beads were then washed 5 times with lysis buffer and resuspended in 2x SDS Laemmli buffer (120 mM Tris-HCl pH 6.8, 4% SDS, 20% glycerol, 0.02% bromophenol blue and 2.5% β-mercatoethanol). 4% of input lysate was loaded unless stated otherwise. Proteins were visualised with InstantBlue stain (Expedeon) or by immunoblotting.

### Immunoblotting

The following primary antibodies were used: anti-XRCC4 (BD611506, BD Biosciences, 1:1,000 and sc-271087, 1:100), anti-mCherry (632543, Takara, 1:1,000), anti-SUMO2/3 (ab3742, Abcam, 1:1,000), anti-GFP (11814460001, Roche, 1:1,000), anti-XLF (ab33499, Abcam, 1:500), anti-LIG4 (ab193353, Abcam, 1:1,000), anti-GST (27457701V, G&E, 1:1000), anti-His (MA1-21315, Invitrogen, 1:1,000). Proteins separated by SDS-PAGE were detected following the manufacturer’s guidelines (GE Healthcare ECL Western Blotting detection system) and the images collected using a Chemidoc imaging system (BioRad Laboratories).

### Recombinant proteins and peptides

Proteins were expressed in *Escherichia coli* BL21-CodonPlus (DE3)-RIL (Stratagene) in Luria-Bertani (LB) medium unless specified otherwise. 6xHis-tagged full-length XRCC4^148^, XRCC4 ^1-213^ (C-to-A: cysteines mutated to alanines)^78^, XRCC4^1-164^ (C-to-A: cysteines mutated to alanines)^104^ expression plasmids were a generous gift from Tom Blundell (University of Cambridge, UK). All XRCC4 recombinant constructs were purified as described previously^78^. The 6xHis-4xSUMO2-Strep expression plasmid was a kind gift from Cynthia Wolberger (Johns Hopkins University, USA)^67^. 6xHis-4xSUMO2-Strep, a linear fusion of a N-terminal full-length SUMO2 fused to truncated SUMO2 (residues 11-92) linked via K11 for the second, third and fourth, was expressed and purified as described previously^67^. pAS2974 (encoding GST-SUMO1)^149^, pAS2976 (encoding GST-SUMO3)^149^ and (encoding GST-4xSUMO2, a linear SUMO2 chain consisting of 4 truncated SUMO2 (residues 12-93) moieties fused to an N-terminal GST tag)^25^ expressing plasmids were kindly provided by Andrew Sharrocks (University of Manchester, UK). GST-SUMO1, GST-SUMO2 and GST-diSUMO2 were purified using glutathione sepharose 4B beads (GE Healthcare) according to the manufacturer’s guidelines, followed by overnight on-bead thrombin digestion (Sigma) and size exclusion chromatography using a Hiload 16/600 Superdex 75 prep grade column (GE Healthcare). GST-4xSUMO2 was purified using MagneGST Glutathione particles (Promega) as previously described^150^. XLF-6xHis was purified using a HP His-trap column followed by a HiTrap Q HP column (both GE Healthcare). mSUMO1 (#UL-712), mSUMO2 (#UL-752), and SUMO2/3 chains enzymatically linked via their internal K11 residues – diSUMO2 (#ULC-200), polySUMO2 (# ULC-210) and polySUMO3 (#ULC-220) – were purchased from Boston Biochem. Recombinant 6xHis-XLF (NBP2-23291), used for SPR experiments, and GST-TCEAL6 (H00158931-P01) were purchased from Novus Biologicals. PAK3-6xHis, 6xHis-WARS (14827-H07B), 6xHis-DARS (14278-H07E), and 6xHis-STMN1 (15440-H07E) were purchased from Sino Biological. 6xHis-SSB (ab84477) and 6xHis-RBBP5 (ab268918) were purchased from Abcam. The peptides used in this study were purchased from Genosphere Biotech. All dialyses were carried out using Spectra/Por 3 kDa MWCO dialysis membranes (Spectrum labs) at 4°C. Proteins were concentrated using Vivaspin PES concentrators (Generon). Protein purity was assessed by protein staining of SDS-PAGE gels.

^15^N-labelled proteins for NMR experiments were expressed in minimal medium supplemented with 15NH_4_Cl. 15N-13C-labelled recombinant XRCC41-180 for NMR assignment was expressed in minimal medium supplemented with 15NH_4_Cl and 13C-glucose. 15N-13C-2H-labelled recombinant XRCC4^1-164^ for NMR assignment was expressed in minimal medium supplemented with 15NH_4_Cl and 13C-glucose and using D_2_O instead of H_2_O for 8 h at 37°C after IPTG induction. Labelled proteins were purified as described above.

## Acknowledgements

We thank Takashi Ochi (University of Leeds, UK) for advice on structural XRCC4 features and critical reading of the manuscript, Tom Blundell (University of Cambridge, UK) for hosting M.J.C.-L. for a short-term visit, Tom Jowitt, Paul Mould and Holly Birchenough (Biomols core facility, UoM) for help with setting up the biophysical binding assays, Siyue Wang for assistance with some of the biochemical procedures, Konstancja Urbaniak for insights into known SUMO receptors, Edward A. McKenzie (MIB) for advice on protein purification, Tom Blundell (University of Cambridge, UK), Andrew Sharrocks (University of Manchester, UK), Cynthia Wolberger (Johns Hopkins University, USA), Edith Heard (EMBL, Heidelberg, Germany), and Murray Junop (Western University London, USA) for kindly providing reagents, and Andrew Blackford (University of Oxford, UK) for critical reading of the manuscript. M.J.C.-L. and C.K.S acknowledge funding by a David Phillips Fellowship BBSRC to C.K.S (BB/N019997/1), and a core research facility pump-priming award (FBMH, UoM) to C.K.S. M.J.C.-L is the recipient of a BBSRC Future Leader Fellowship (BB/R01212/1). This research was in part funded by the Wellcome Trust. S.P.J. acknowledges funding by: Wellcome Investigator Award 206388/Z/17/Z, Cancer Research UK Programme Grant C6/A18796, ERC Synergy Grant (DDREAMM), ERC Advanced Researcher Grant, GA 268536 (DDRREAM), and Gurdon Institute core infrastructure funding by Cancer Research UK C6946/A24843 and Wellcome WT203144. Q.W. is a University Academic Fellow at the University of Leeds. The Bruker MaXis II instrument used in this study was funded by the BBSRC (BB/M017982/1). For the purpose of open access, the author has applied a CC BY public copyright licence to any author accepted manuscript version arising from this submission.

## References

1. Flotho, A. & Melchior, F. Sumoylation: a regulatory protein modification in health and disease. Annu. Rev. Biochem. 82, 357–85 (2013).

2. Pichler, A., Fatouros, C., Lee, H. & Eisenhardt, N. SUMO conjugation - A mechanistic view. Biomol. Concepts 8, 13–36 (2017).

3. Cubeñas-Potts, C. & Matunis, M. J. SUMO: A Multifaceted Modifier of Chromatin Structure and Function. Dev. Cell 24, 1–12 (2013).

4. Barry, J. & Lock, R. B. Small ubiquitin-related modifier-1: Wrestling with protein regulation. Int. J. Biochem. Cell Biol. 43, 37–40 (2011).

5. Hay, R. T. Decoding the SUMO signal. Biochem. Soc. Trans. 41, 463–473 (2013).

6. Droescher, M., Chaugule, V. K. & Pichler, A. SUMO rules: Regulatory concepts and their implication in neurologic functions. NeuroMolecular Med. 15, 639–660 (2013).

7. Shetty, P. M. V., Rangrez, A. Y. & Frey, N. SUMO proteins in the cardiovascular system: friend or foe? J. Biomed. Sci. 27, 98 (2020).

8. Kroonen, J. S. & Vertegaal, A. C. O. Targeting SUMO Signaling to Wrestle Cancer. Trends in Cancer (2021) doi:10.1016/j.trecan.2020.11.009.

9. Vertegaal, A. C. O. SUMO chains: polymeric signals. Biochem. Soc. Trans. 38, 46–49 (2010).

10. Ulrich, H. D. The Fast-Growing Business of SUMO Chains. Mol. Cell 32, 301–305 (2008).

11. Jansen, N. S. & Vertegaal, A. C. O. A Chain of Events: Regulating Target Proteins by SUMO Polymers. Trends Biochem. Sci. (2021) doi:10.1016/j.tibs.2020.09.002.

12. Hendriks, I. A. & Vertegaal, A. C. O. A comprehensive compilation of SUMO proteomics. Nat. Rev. Mol. Cell Biol. 17, 581–595 (2016).

13. Beauclair, G., Bridier-Nahmias, A., Zagury, J.-F., Saïb, A. & Zamborlini, A. JASSA: a comprehensive tool for prediction of SUMOylation sites and SIMs. Bioinformatics 31, 3483–3491 (2015).

14. Husnjak, K. & Dikic, I. Ubiquitin-Binding Proteins: Decoders of Ubiquitin-Mediated Cellular Functions. Annu. Rev. Biochem. 81, 291–322 (2012).

15. Praefcke, G. J. K., Hofmann, K. & Dohmen, R. J. SUMO playing tag with ubiquitin. Trends Biochem. Sci. 37, 23–31 (2012).

16. Hecker, C.-M., Rabiller, M., Haglund, K., Bayer, P. & Dikic, I. Specification of SUMO1- and SUMO2-interacting Motifs. J. Biol. Chem. 281, 16117–16127 (2006).

17. Sun, H. & Hunter, T. Poly-Small Ubiquitin-like Modifier (PolySUMO)-binding Proteins Identified through a String Search. J. Biol. Chem. 287, 42071–42083 (2012).

18. Minty, A., Dumont, X., Kaghad, M. & Caput, D. Covalent Modification of p73α by SUMO-1. J. Biol. Chem. 275, 36316–36323 (2000).

19. Song, J., Durrin, L. K., Wilkinson, T. A., Krontiris, T. G. & Chen, Y. Identification of a SUMO-binding motif that recognizes SUMO-modified proteins. Proc. Natl. Acad. Sci. U. S. A. 101, 14373–14378 (2004).

20. Zhao, B., Rothenberg, E., Ramsden, D. A. & Lieber, M. R. The molecular basis and disease relevance of non-homologous DNA end joining. Nat. Rev. Mol. Cell Biol. (2020) doi:10.1038/s41580-020-00297-8.

21. Jackson, S. P. & Bartek, J. The DNA-damage response in human biology and disease. Nature 461, 1071–1078 (2009).

22. Ciccia, A. & Elledge, S. J. The DNA damage response: making it safe to play with knives. Mol. Cell 40, 179–204 (2010).

23. Schwertman, P., Bekker-Jensen, S. & Mailand, N. Regulation of DNA double-strand break repair by ubiquitin and ubiquitin-like modifiers. Nat. Rev. Mol. Cell Biol. 17, 379–394 (2016).

24. Garvin, A. J. & Morris, J. R. SUMO, a small, but powerful, regulator of double-strand break repair. Philos. Trans. R. Soc. B Biol. Sci. 372, 20160281 (2017).

25. Aguilar-Martinez, E. et al. Screen for multi-SUMO–binding proteins reveals a multi-SIM–binding mechanism for recruitment of the transcriptional regulator ZMYM2 to chromatin. Proc. Natl. Acad. Sci. 112, E4854–E4863 (2015).

26. Makhnevych, T. et al. Global Map of SUMO Function Revealed by Protein-Protein Interaction and Genetic Networks. Mol. Cell 33, 124–135 (2009).

27. Ouyang, J., Shi, Y., Valin, A., Xuan, Y. & Gill, G. Direct Binding of CoREST1 to SUMO-2/3 Contributes to Gene-Specific Repression by the LSD1/CoREST1/HDAC Complex. Mol. Cell 34, 145–154 (2009).

28. Hannich, J. T. et al. Defining the SUMO-modified Proteome by Multiple Approaches in Saccharomyces cerevisiae. J. Biol. Chem. 280, 4102–4110 (2005).

29. Rosendorff, A. et al. NXP-2 association with SUMO-2 depends on lysines required for transcriptional repression. Proc. Natl. Acad. Sci. 103, 5308 LP – 5313 (2006).

30. Desterro, J. M. P., Rodriguez, M. S., Kemp, G. D. & Hay, R. T. Identification of the Enzyme Required for Activation of the Small Ubiquitin-like Protein SUMO-1. J. Biol. Chem. 274, 10618–10624 (1999).

31. Gong, L., Li, B., Millas, S. & Yeh, E. T. H. Molecular cloning and characterization of human AOS1 and UBA2, components of the sentrin-activating enzyme complex. FEBS Lett. 448, 185–189 (1999).

32. Okuma, T., Honda, R., Ichikawa, G., Tsumagari, N. & Yasuda, H. In VitroSUMO-1 Modification Requires Two Enzymatic Steps, E1 and E2. Biochem. Biophys. Res. Commun. 254, 693–698 (1999).

33. Olsen, S. K., Capili, A. D., Lu, X., Tan, D. S. & Lima, C. D. Active site remodelling accompanies thioester bond formation in the SUMO E1. Nature 463, 906–912 (2010).

34. Lin, D.-Y. et al. Role of SUMO-Interacting Motif in Daxx SUMO Modification, Subnuclear Localization, and Repression of Sumoylated Transcription Factors. Mol. Cell 24, 341–354 (2006).

35. Chang, C.-C. et al. Structural and Functional Roles of Daxx SIM Phosphorylation in SUMO Paralog-Selective Binding and Apoptosis Modulation. Mol. Cell 42, 62–74 (2011).

36. Escobar-Cabrera, E. et al. Characterizing the N- and C-terminal Small Ubiquitin-like Modifier (SUMO)-interacting Motifs of the Scaffold Protein DAXX. J. Biol. Chem. 286, 19816–19829 (2011).

37. Widagdo, J., Taylor, K. M., Gunning, P. W., Hardeman, E. C. & Palmer, S. J. SUMOylation of GTF2IRD1 Regulates Protein Partner Interactions and Ubiquitin-Mediated Degradation. PLoS One 7, e49283 (2012).

38. Fattet, L. et al. TIF1γ requires sumoylation to exert its repressive activity on TGFβ signaling. J. Cell Sci. 126, 3713 LP – 3723 (2013).

39. Lecona, E. et al. USP7 is a SUMO deubiquitinase essential for DNA replication. Nat. Struct. Mol. Biol. 23, 270–277 (2016).

40. Guzzo, C. M. & Matunis, M. J. Expanding SUMO and ubiquitin-mediated signaling through hybrid SUMO-ubiquitin chains and their receptors. Cell Cycle 12, 1015–1017 (2013).

41. Brüninghoff, K. et al. Identification of SUMO binding proteins enriched after covalent photo-crosslinking. ACS Chem. Biol. (2020) doi:10.1021/acschembio.0c00609.

42. Garzón, J. et al. SUMO-SIM Interactions Regulate the Activity of RGSZ2 Proteins. PLoS One 6, e28557 (2011).

43. Pilla, E. et al. A Novel SUMO1-specific Interacting Motif in Dipeptidyl Peptidase 9 (DPP9) That Is Important for Enzymatic Regulation. J. Biol. Chem. 287, 44320–44329 (2012).

44. Um, J. W. & Chung, K. C. Functional modulation of parkin through physical interaction with SUMO-1. J. Neurosci. Res. 84, 1543–1554 (2006).

45. Brocchieri, L. & Karlin, S. Protein length in eukaryotic and prokaryotic proteomes. Nucleic Acids Res. 33, 3390–3400 (2005).

46. Danielsen, J. R. et al. DNA damage–inducible SUMOylation of HERC2 promotes RNF8 binding via a novel SUMO-binding Zinc finger. J. Cell Biol. 197, 179–187 (2012).

47. Zhang, X.-D. et al. SUMO-2/3 Modification and Binding Regulate the Association of CENP-E with Kinetochores and Progression through Mitosis. Mol. Cell 29, 729–741 (2008).

48. Diehl, C. et al. Structural Analysis of a Complex between Small Ubiquitin-like Modifier 1 (SUMO1) and the ZZ Domain of CREB-binding Protein (CBP/p300) Reveals a New Interaction Surface on SUMO. J. Biol. Chem. 291, 12658–12672 (2016).

49. Goodarzi, A. A., Kurka, T. & Jeggo, P. A. KAP-1 phosphorylation regulates CHD3 nucleosome remodeling during the DNA double-strand break response. Nat. Struct. Mol. Biol. 18, 831–839 (2011).

50. Ouyang, J. et al. Noncovalent Interactions with SUMO and Ubiquitin Orchestrate Distinct Functions of the SLX4 Complex in Genome Maintenance. Mol. Cell 57, 108–122 (2015).

51. Enserink, J. M. Sumo and the cellular stress response. Cell Div. 10, 4 (2015).

52. Saitoh, H. & Hinchey, J. Functional Heterogeneity of Small Ubiquitin-related Protein Modifiers SUMO-1 versus SUMO-2/3. J. Biol. Chem. 275, 6252–6258 (2000).

53. Morris, J. R. & Garvin, A. J. SUMO in the DNA Double-Stranded Break Response: Similarities, Differences, and Cooperation with Ubiquitin. J. Mol. Biol. 429, 3376–3387 (2017).

54. Ulrich, H. D. SUMO Teams up with Ubiquitin to Manage Hypoxia. Cell 131, 446–447 (2007).

55. Cox, E. et al. Global Analysis of SUMO-Binding Proteins Identifies SUMOylation as a Key Regulator of the INO80 Chromatin Remodeling Complex. Mol. Cell. Proteomics 16, 812–823 (2017).

56. Keiten-Schmitz, J., Schunck, K. & Müller, S. SUMO Chains Rule on Chromatin Occupancy. Frontiers in Cell and Developmental Biology vol. 7 343 (2020).

57. Wasik, U. & Filipek, A. Non-nuclear function of sumoylated proteins. Biochim. Biophys. Acta - Mol. Cell Res. 1843, 2878–2885 (2014).

58. Alonso, A. et al. Emerging roles of sumoylation in the regulation of actin, microtubules, intermediate filaments, and septins. Cytoskeleton 72, 305–339 (2015).

59. Hendriks, I. A. et al. Site-specific mapping of the human SUMO proteome reveals co-modification with phosphorylation. Nat. Struct. Mol. Biol. 24, 325–336 (2017).

60. Tomanov, K. et al. Sumoylation and phosphorylation: hidden and overt links. J. Exp. Bot. 69, 4583–4590 (2018).

61. Tang, J. et al. Critical role for Daxx in regulating Mdm2. Nat. Cell Biol. 8, 855–862 (2006).

62. Nie, M. et al. Dual recruitment of Cdc48 (p97)-Ufd1-Npl4 ubiquitin-selective segregase by small ubiquitin-like modifier protein (SUMO) and ubiquitin in SUMO-targeted ubiquitin ligase-mediated genome stability functions. J. Biol. Chem. 287, 29610–29619 (2012).

63. Chanarat, S. & Mishra, S. K. Emerging Roles of Ubiquitin-like Proteins in Pre-mRNA Splicing. Trends Biochem. Sci. 43, 896–907 (2018).

64. Pozzi, B., Mammi, P., Bragado, L., Giono, L. E. & Srebrow, A. When SUMO met splicing. RNA Biol. 15, 689–695 (2018).

65. Chymkowitch, P. et al. TORC1-dependent sumoylation of Rpc82 promotes RNA polymerase III assembly and activity. Proc. Natl. Acad. Sci. 114, 1039–1044 (2017).

66. Nguéa P, A., Robertson, J., Herrera, M. C., Chymkowitch, P. & Enserink, J. M. Desumoylation of RNA polymerase III lies at the core of the Sumo stress response in yeast. J. Biol. Chem. 294, 18784–18795 (2019).

67. DiBello, A., Datta, A. B., Zhang, X. & Wolberger, C. Role of E2-RING Interactions in Governing RNF4-Mediated Substrate Ubiquitination. J. Mol. Biol. 428, 4639–4650 (2016).

68. Xu, Y. et al. Structural insight into SUMO chain recognition and manipulation by the ubiquitin ligase RNF4. Nat. Commun. 5, 4217 (2014).

69. Kost, L. J. & Mootz, H. D. A FRET Sensor to Monitor Bivalent SUMO–SIM Interactions in SUMO Chain Binding. ChemBioChem 19, 177–184 (2018).

70. Sriramachandran, A. M. et al. Arkadia/RNF111 is a SUMO-targeted ubiquitin ligase with preference for substrates marked with SUMO1-capped SUMO2/3 chain. Nat. Commun. 10, 3678 (2019).

71. Bouchenna, J., Sénéchal, M., Drobecq, H., Vicogne, J. & Melnyk, O. Total Chemical Synthesis of All SUMO-2/3 Dimer Combinations. Bioconjug. Chem. 30, 2967–2973 (2019).

72. Cuijpers, S. A. G. et al. Chromokinesin KIF4A teams up with stathmin 1 to regulate abscission in a SUMO-dependent manner. J. Cell Sci. 133, jcs248591 (2020).

73. Zhao, Q. et al. GPS-SUMO: a tool for the prediction of sumoylation sites and SUMO-interaction motifs. Nucleic Acids Res. 42, W325–W330 (2014).

74. Galanty, Y. et al. Mammalian SUMO E3-ligases PIAS1 and PIAS4 promote responses to DNA double-strand breaks. Nature 462, 935–9 (2009).

75. Galanty, Y., Belotserkovskaya, R., Coates, J. & Jackson, S. P. RNF4, a SUMO-targeted ubiquitin E3 ligase, promotes DNA double-strand break repair. Genes Dev. 26, 1179–1195 (2012).

76. Li, Y.-J., Stark, J. M., Chen, D. J., Ann, D. K. & Chen, Y. Role of SUMO:SIM-mediated protein-protein interaction in non-homologous end joining. Oncogene 29, 3509–3518 (2010).

77. Junop, M. S. Crystal structure of the Xrcc4 DNA repair protein and implications for end joining. EMBO J. 19, (2000).

78. Sibanda, B. L. Crystal structure of an Xrcc4–DNA ligase IV complex. Nat. Struct. Biol. 8, (2001).

79. Yurchenko, V., Xue, Z. & Sadofsky, M. J. SUMO Modification of Human XRCC4 Regulates Its Localization and Function in DNA Double-Strand Break Repair. Mol. Cell. Biol. 26, 1786 LP – 1794 (2006).

80. Hammel, M., Yu, Y., Fang, S., Lees-Miller, S. P. & Tainer, J. A. XLF Regulates Filament Architecture of the XRCC4·Ligase IV Complex. Structure 18, 1431–1442 (2010).

81. Li, W. et al. The nucleoskeleton protein IFFO1 immobilizes broken DNA and suppresses chromosome translocation during tumorigenesis. Nat. Cell Biol. 21, 1273–1285 (2019).

82. Wu, P. Y. Structural and functional interaction between the human DNA repair proteins DNA ligase IV and XRCC4. Mol. Cell. Biol. 29, (2009).

83. Hammel, M. XRCC4 protein interactions with XRCC4-like factor (XLF) create an extended grooved scaffold for DNA ligation and double strand break repair. J. Biol. Chem. 286, (2011).

84. Aceytuno, R. D. et al. Structural and functional characterization of the PNKP–XRCC4–LigIV DNA repair complex. Nucleic Acids Res. 45, 6238–6251 (2017).

85. Hammel, M. et al. An Intrinsically Disordered APLF Links Ku, DNA-PKcs, and XRCC4-DNA Ligase IV in an Extended Flexible Non-homologous End Joining Complex. J. Biol. Chem. 291, 26987–27006 (2016).

86. Clements, P. M. et al. The ataxia–oculomotor apraxia 1 gene product has a role distinct from ATM and interacts with the DNA strand break repair proteins XRCC1 and XRCC4. DNA Repair (Amst). 3, 1493–1502 (2004).

87. Cherry, A. L. et al. Versatility in phospho-dependent molecular recognition of the XRCC1 and XRCC4 DNA-damage scaffolds by aprataxin-family FHA domains. DNA Repair (Amst). 35, 116–125 (2015).

88. Goodarzi, A. A. et al. DNA-PK autophosphorylation facilitates Artemis endonuclease activity. EMBO J. 25, 3880–3889 (2006).

89. Jette, N. & Lees-Miller, S. P. The DNA-dependent protein kinase: A multifunctional protein kinase with roles in DNA double strand break repair and mitosis. Prog. Biophys. Mol. Biol. 117, 194–205 (2015).

90. Andres, S. N. et al. A human XRCC4–XLF complex bridges DNA. Nucleic Acids Res. 40, 1868–1878 (2012).

91. Ahnesorg, P., Smith, P. & Jackson, S. P. XLF Interacts with the XRCC4-DNA Ligase IV Complex to Promote DNA Nonhomologous End-Joining. Cell 124, 301–313 (2006).

92. Roy, S. et al. XRCC4/XLF Interaction Is Variably Required for DNA Repair and Is Not Required for Ligase IV Stimulation. Mol. Cell. Biol. 35, 3017–3028 (2015).

93. Graham, T. G. W., Walter, J. C. & Loparo, J. J. Two-Stage Synapsis of DNA Ends during Non-homologous End Joining. Mol. Cell 61, 850–858 (2016).

94. Zhao, B. et al. The essential elements for the noncovalent association of two DNA ends during NHEJ synapsis. Nat. Commun. 10, 3588 (2019).

95. Graham, T. G. W., Carney, S. M., Walter, J. C. & Loparo, J. J. A single XLF dimer bridges DNA ends during nonhomologous end joining. Nat. Struct. Mol. Biol. 25, 877–884 (2018).

96. Nick McElhinny, S. A., Snowden, C. M., McCarville, J. & Ramsden, D. A. Ku Recruits the XRCC4-Ligase IV Complex to DNA Ends. Mol. Cell. Biol. 20, 2996–3003 (2000).

97. Srikumar, T. et al. Global analysis of SUMO chain function reveals multiple roles in chromatin regulation. J. Cell Biol. 201, 145–163 (2013).

98. Tatham, M. H. et al. Polymeric Chains of SUMO-2 and SUMO-3 Are Conjugated to Protein Substrates by SAE1/SAE2 and Ubc9. J. Biol. Chem. 276, 35368–35374 (2001).

99. Bruderer, R. et al. Purification and identification of endogenous polySUMO conjugates. EMBO Rep. 12, 142–148 (2011).

100. Matic, I. et al. In vivo identification of human small ubiquitin-like modifier polymerization sites by high accuracy mass spectrometry and an in vitro to in vivo strategy. Mol. Cell. Proteomics 7, 132–144 (2008).

101. Normanno, D. et al. Mutational phospho-mimicry reveals a regulatory role for the XRCC4 and XLF C-terminal tails in modulating DNA bridging during classical non-homologous end joining. Elife 6, (2017).

102. Kerscher, O. SUMO junction-what’s your function? New insights through SUMO-interacting motifs. EMBO Rep 8, 550–555 (2007).

103. Ropars, V. Structural characterization of filaments formed by human Xrcc4–Cernunnos/XLF complex involved in nonhomologous DNA end-joining. Proc. Natl Acad. Sci. USA 108, (2011).

104. Wu, Q. et al. Non-homologous end-joining partners in a helical dance: structural studies of XLF–XRCC4 interactions. Biochem. Soc. Trans. 39, 1387–1392 (2011).

105. Manzi, L. et al. Carbene footprinting accurately maps binding sites in protein–ligand and protein–protein interactions. Nat. Commun. 7, 13288 (2016).

106. Jenner, M. et al. Mechanism of intersubunit ketosynthase–dehydratase interaction in polyketide synthases. Nat. Chem. Biol. 14, 270–275 (2018).

107. Manzi, L. et al. Carbene Footprinting Reveals Binding Interfaces of a Multimeric Membrane-Spanning Protein. Angew. Chemie Int. Ed. 56, 14873–14877 (2017).

108. Song, J., Zhang, Z., Hu, W. & Chen, Y. Small ubiquitin-like modifier (SUMO) recognition of a SUMO binding motif: a reversal of the bound orientation. J. Biol. Chem. 280, 40122–9 (2005).

109. Andres, S. N., Modesti, M., Tsai, C. J., Chu, G. & Junop, M. S. Crystal Structure of Human XLF: A Twist in Nonhomologous DNA End-Joining. Mol. Cell 28, 1093–1101 (2007).

110. Jackson, S. P. & Durocher, D. Regulation of DNA Damage Responses by Ubiquitin and SUMO. Mol. Cell 49, 795–807 (2013).

111. Schellenberg, M. J. et al. ZATT (ZNF451)–mediated resolution of topoisomerase 2 DNA-protein cross-links. Science (80-.). 357, 1412–1416 (2017).

112. Rulten, S. L. & Grundy, G. J. Non-homologous end joining: Common interaction sites and exchange of multiple factors in the DNA repair process. BioEssays 39, 1600209 (2017).

113. Kumar, V., Alt, F. W. & Frock, R. L. PAXX and XLF DNA repair factors are functionally redundant in joining DNA breaks in a G1-arrested progenitor B-cell line. Proc. Natl. Acad. Sci. U. S. A. 113, 10619–10624 (2016).

114. Hung, P. J. et al. MRI Is a DNA Damage Response Adaptor during Classical Non-homologous End Joining. Mol. Cell 71, 332–342.e8 (2018).

115. Zha, S. et al. ATM damage response and XLF repair factor are functionally redundant in joining DNA breaks. Nature 469, 250–254 (2011).

116. Beck, C., Castañeda-Zegarra, S., Huse, C., Xing, M. & Oksenych, V. Mediator of DNA Damage Checkpoint Protein 1 Facilitates V(D)J Recombination in Cells Lacking DNA Repair Factor XLF. Biomolecules 10, 60 (2019).

117. Liu, X. et al. Overlapping functions between XLF repair protein and 53BP1 DNA damage response factor in end joining and lymphocyte development. Proc. Natl. Acad. Sci. 109, 3903 LP – 3908 (2012).

118. Brown, J. S. et al. Neddylation promotes ubiquitylation and release of Ku from DNA-damage sites. Cell Rep. 11, 704–714 (2015).

119. Postow, L. et al. Ku80 removal from DNA through double strand break-induced ubiquitylation. J. Cell Biol. 182, 467–479 (2008).

120. Postow, L. Destroying the ring: Freeing DNA from Ku with ubiquitin. FEBS Lett. 585, 2876–2882 (2011).

121. van den Boom, J. et al. VCP/p97 Extracts Sterically Trapped Ku70/80 Rings from DNA in Double-Strand Break Repair. Mol. Cell 64, 189–198 (2016).

122. Liu, P. et al. Akt-mediated phosphorylation of XLF impairs non-homologous end-joining DNA repair. Mol. Cell 57, 648–661 (2015).

123. Hentges, P., Waller, H., Reis, C. C., Ferreira, M. G. & Doherty, A. J. Cdk1 restrains NHEJ through phosphorylation of XRCC4-like factor Xlf1. Cell Rep. 9, 2011–2017 (2014).

124. Ray Chaudhuri, A. & Nussenzweig, A. The multifaceted roles of PARP1 in DNA repair and chromatin remodelling. Nat. Rev. Mol. Cell Biol. 18, 610–621 (2017).

125. Osborne, H. C., Irving, E. & Schmidt, C. K. The Ubiquitin/UBL Drug Target Repertoire. Trends Mol. Med. 26, 1133–1134 (2020).

126. Menon, V. & Povirk, L. F. XLF/Cernunnos: An important but puzzling participant in the nonhomologous end joining DNA repair pathway. DNA Repair (Amst). 58, 29–37 (2017).

127. Menchon, G. et al. Structure-Based Virtual Ligand Screening on the XRCC4/DNA Ligase IV Interface. Sci. Rep. 6, 22878 (2016).

128. Canny, M. D. et al. Inhibition of 53BP1 favors homology-dependent DNA repair and increases CRISPR-Cas9 genome-editing efficiency. Nat. Biotechnol. 36, 95–102 (2018).

129. Mi, H., Muruganujan, A., Casagrande, J. T. & Thomas, P. D. Large-scale gene function analysis with the PANTHER classification system. Nat. Protoc. 8, 1551–1566 (2013).

130. Mi, H. et al. Protocol Update for large-scale genome and gene function analysis with the PANTHER classification system (v.14.0). Nat. Protoc. 14, 703–721 (2019).

131. Szklarczyk, D. et al. STRING v11: protein–protein association networks with increased coverage, supporting functional discovery in genome-wide experimental datasets. Nucleic Acids Res. 47, D607–D613 (2019).

132. Naik, M. T., Naik, N., Shih, H., Huang, T. Solution structure of human SUMO2. (2016) doi:10.2210/pdb2N1W/pdb.

133. Baba, D. et al. Crystal Structure of SUMO-3-modified Thymine-DNA Glycosylase. J. Mol. Biol. 359, 137–147 (2006).

134. Jurrus, E. et al. Improvements to the APBS biomolecular solvation software suite. Protein Sci. 27, 112–128 (2018).

135. Krystkowiak, I., Manguy, J. & Davey, N. E. PSSMSearch: a server for modeling, visualization, proteome-wide discovery and annotation of protein motif specificity determinants. Nucleic Acids Res. 46, W235–W241 (2018).

136. Sievers, F. et al. Fast, scalable generation of high-quality protein multiple sequence alignments using Clustal Omega. Mol. Syst. Biol. 7, 539 (2011).

137. Naik, M.T., Naik, N., Shih, H., Huang, T. Solution structure of human SUMO1. doi:10.2210/pdb2N1V/pdb.

138. Ding, H. et al. Solution Structure of Human SUMO-3 C47S and Its Binding Surface for Ubc9,. Biochemistry 44, 2790–2799 (2005).

139. Smyth, G. K. & Speed, T. Normalization of cDNA microarray data. Methods 31, 265–273 (2003).

140. Gardner, K. H. & Kay, L. E. The use of 2 H, 13 C, 15 N multidimensional NMR to study the structure and dynamics of proteins. Annu. Rev. Biophys. Biomol. Struct. 27, 357–406 (1998).

141. Hyberts, S. G., Takeuchi, K. & Wagner, G. Poisson-gap sampling and forward maximum entropy reconstruction for enhancing the resolution and sensitivity of protein NMR data. J. Am. Chem. Soc. 132, 2145—2147 (2010).

142. Shen, Y., Delaglio, F., Cornilescu, G. & Bax, A. TALOS+: a hybrid method for predicting protein backbone torsion angles from NMR chemical shifts. J. Biomol. NMR 44, 213–223 (2009).

143. Vranken, W. F. et al. The CCPN data model for NMR spectroscopy: Development of a software pipeline. Proteins Struct. Funct. Bioinforma. 59, 687–696 (2005).

144. Skinner, S. P. et al. CcpNmr AnalysisAssign: a flexible platform for integrated NMR analysis. J. Biomol. NMR 66, 111–124 (2016).

145. Dominguez, C., Boelens, R. & Bonvin, A. M. J. J. HADDOCK: A Protein−Protein Docking Approach Based on Biochemical or Biophysical Information. J. Am. Chem. Soc. 125, 1731–1737 (2003).

146. Britton, S., Coates, J. & Jackson, S. P. A new method for high-resolution imaging of Ku foci to decipher mechanisms of DNA double-strand break repair. J. Cell Biol. 202, 579–595 (2013).

147. Masui, O. et al. Live-Cell Chromosome Dynamics and Outcome of X Chromosome Pairing Events during ES Cell Differentiation. Cell 145, 447–458 (2011).

148. Wang, J. L. et al. Dissection of DNA double-strand-break repair using novel single-molecule forceps. Nat. Struct. Mol. Biol. 25, 482–487 (2018).

149. Yang, S. & Sharrocks, A. D. The SUMO E3 Ligase Activity of Pc2 Is Coordinated through a SUMO Interaction Motif. Mol. Cell. Biol. 30, 2193 LP – 2205 (2010).

150. Aguilar-Martínez, E. & Sharrocks, A. D. The Use of Multimeric Protein Scaffolds for Identifying Multi-SUMO Binding Proteins BT - SUMO: Methods and Protocols. in (ed. Rodriguez, M. S.) 195–204 (Springer New York, 2016). doi:10.1007/978-1-4939-6358-4_14.

