## Supplementary Figures for "Microarray screening reveals a non-conventional SUMO-binding mode linked to DNA repair by non-homologous end-joining"

### Figure S1

Protein stains

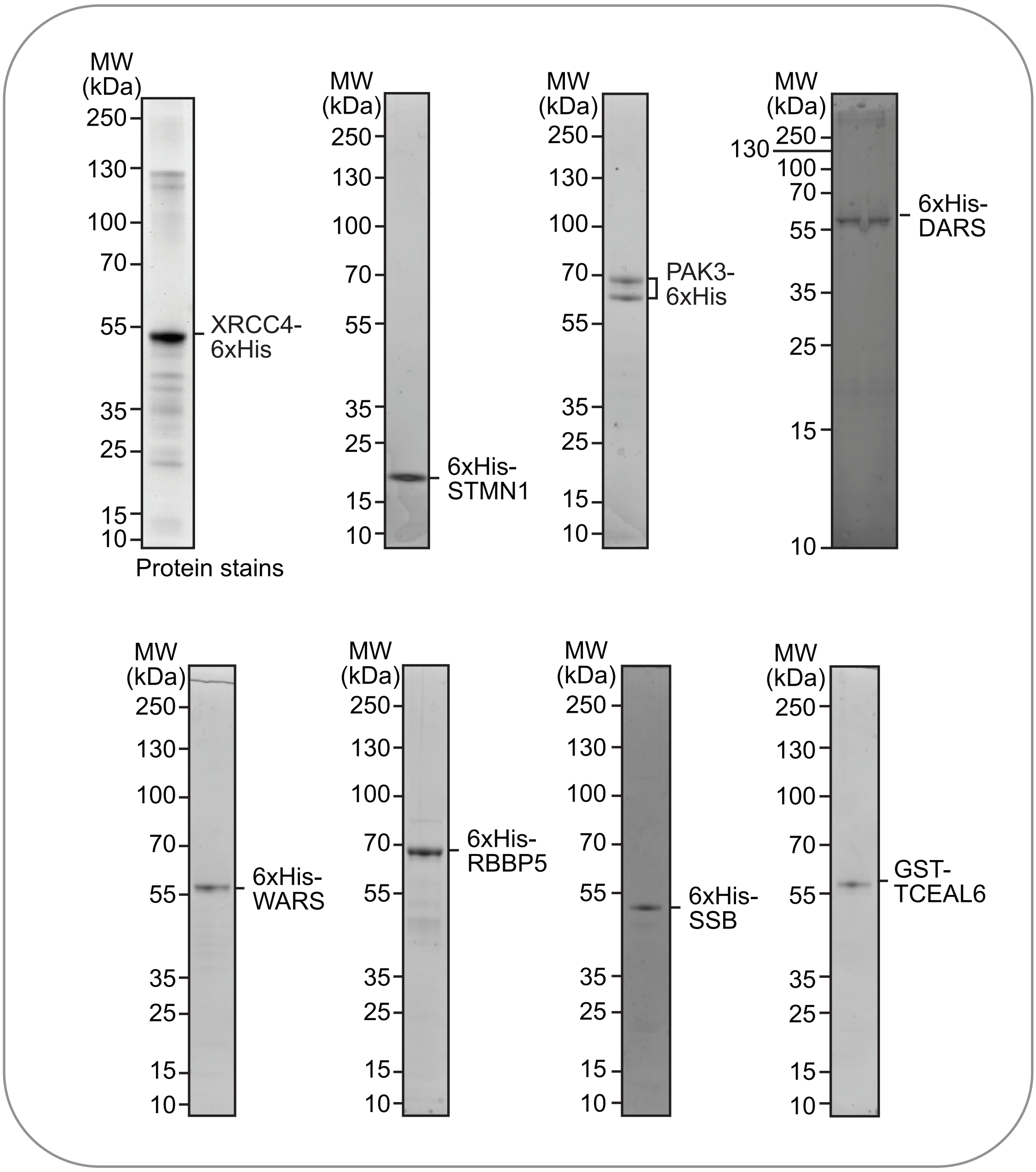

**Figure S2**

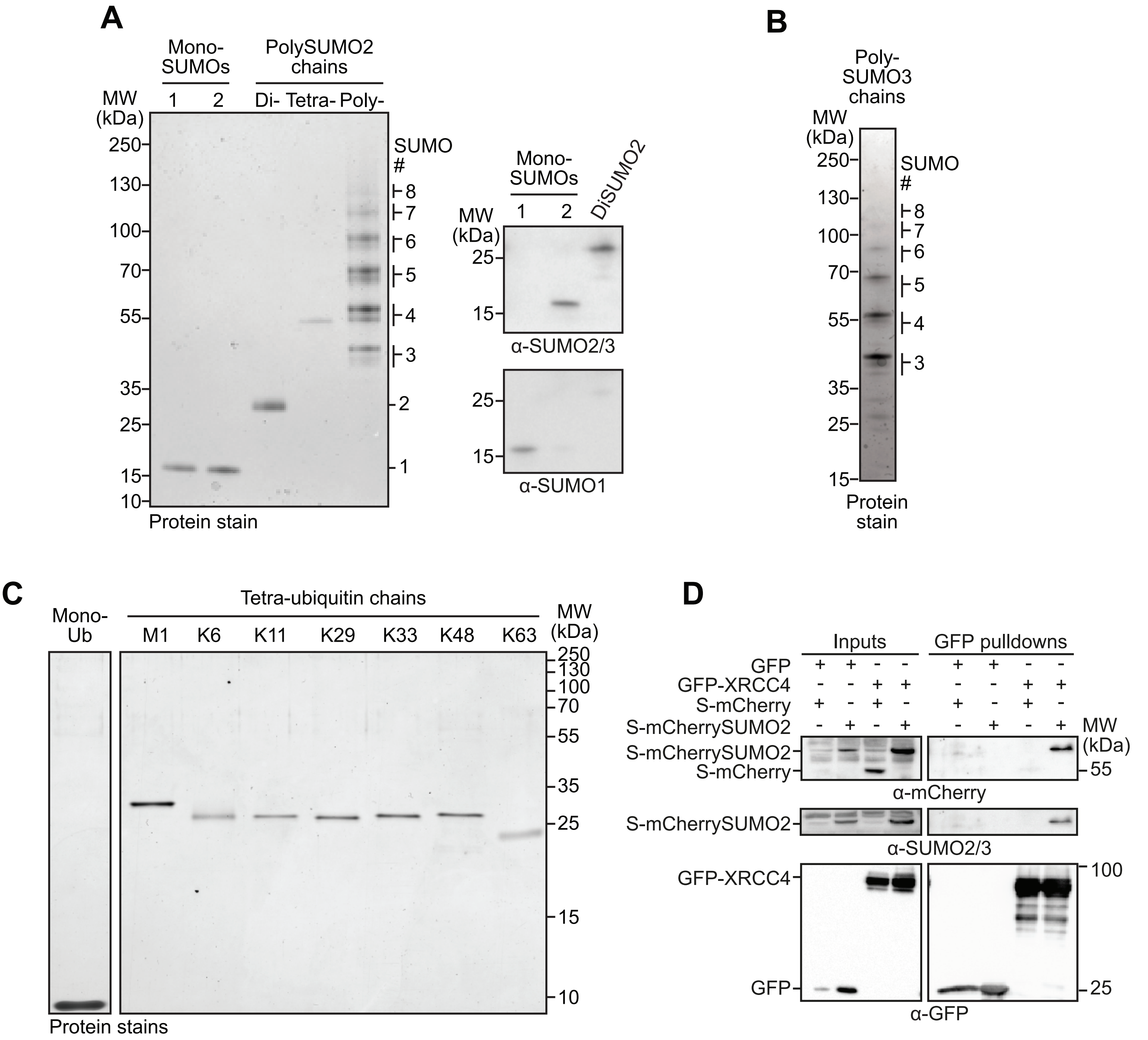
A
Mono-SUMOs
PolySUMO2 chains
MW (kDa)
1
2
Di-
Tetra-
Poly-
250
130
100
70
55
35
25
15
10
SUMO #
8
7
6
5
4
3
2
1
Protein stain
Mono-SUMOs
DiSUMO2
α-SUMO2/3
α-SUMO1
B
Poly-SUMO3 chains
Protein stain
C
Mono-Ub
Tetra-ubiquitin chains
M1
K6
K11
K29
K33
K48
K63
Protein stains
D
Inputs
GFP pulldowns
GFP
GFP-XRCC4
S-mCherry
S-mCherrySUMO2
α-mCherry
α-SUMO2/3
α-GFP

### Figure S3

A

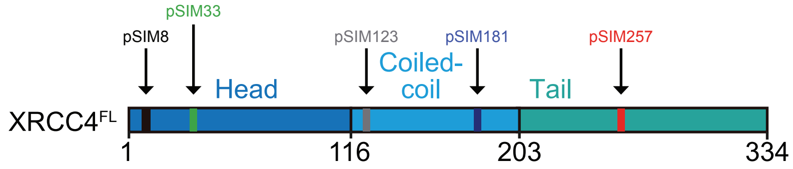


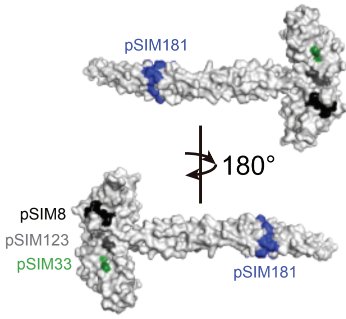


| Putative SIM (GPS-SUMO) | Position | Sequence | | |
|---|---|---|---|---|
| pSIM8 | 8-12 | MERKISR | IHLVS | EPSITHF |
| pSIM33 | 33-37 | KTLESGF | VITLT | DGHSAWT |
| pSIM123 | 123-127 | VENPAEV | IRELI | CYCLDTI |
| pSIM181 | 181-185 | TDLYKRF | ILVLN | EKKTKIR |
| pSIM257 | 257-261 | AVSKDDS | IISSL | DVTDIAP |

B

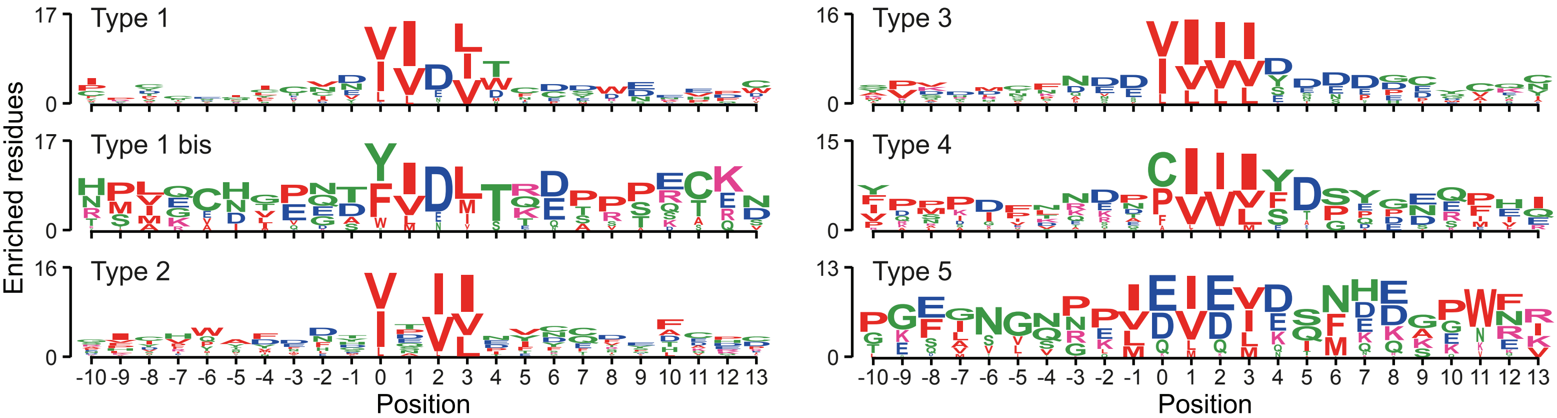


C

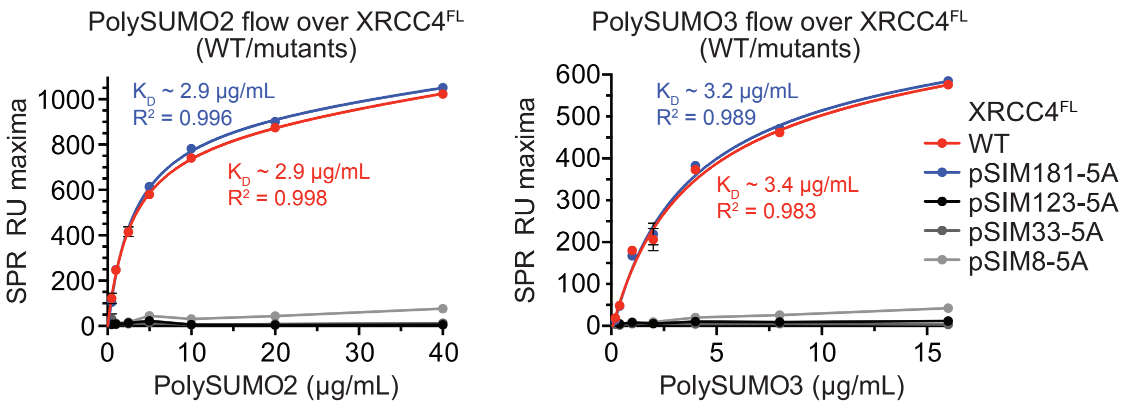


D

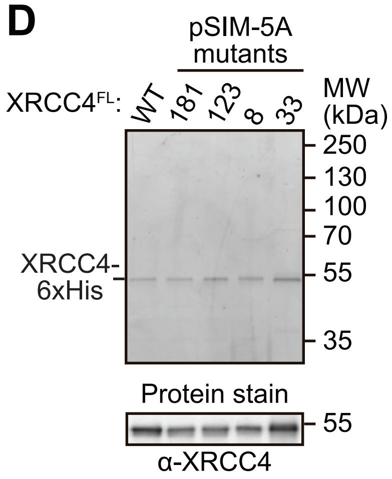


E

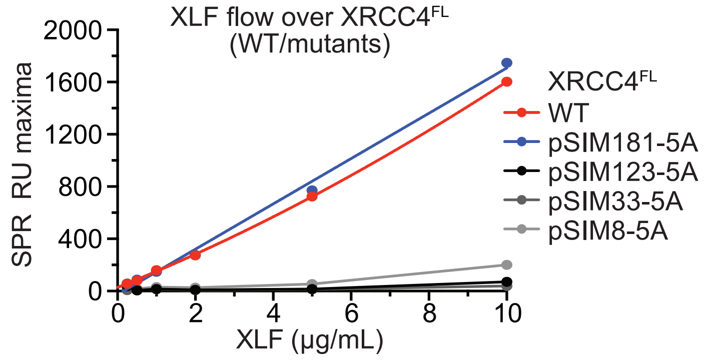


F

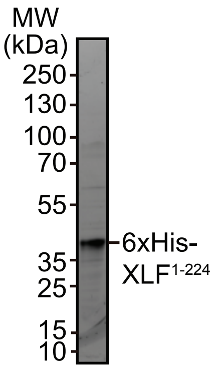


G

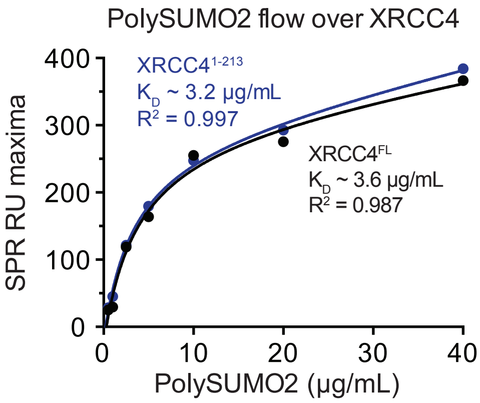

### Figure S4

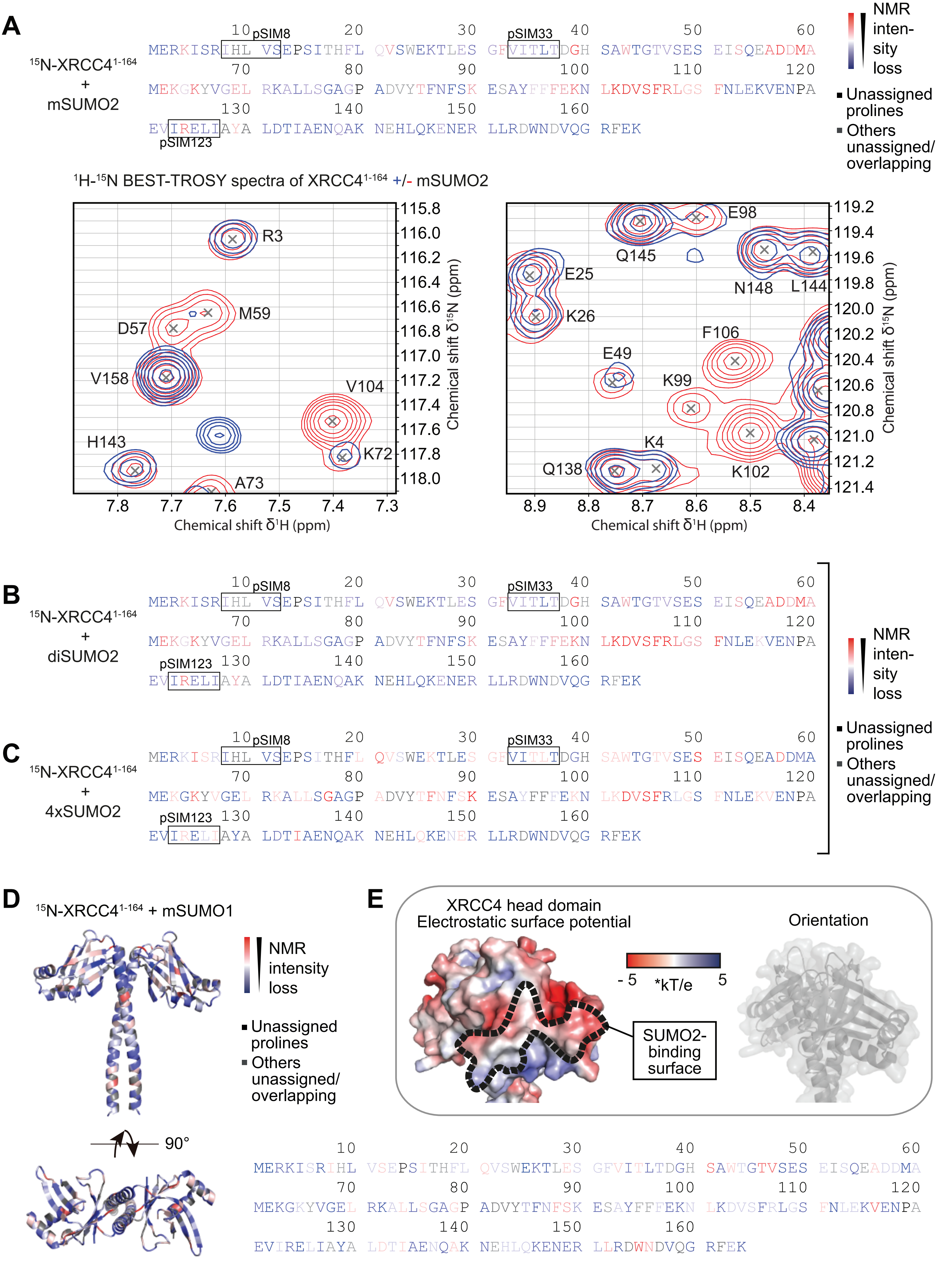
A
15N-XRCC4 1-164 + mSUMO2
MERKISRIHL VSEPSITHFL QVSWEKTLES GFVITLTDGH SAWTGTVSES EISQEADDMA
MEKGKYVGEL RKALLSGAGP ADVYTFNFSK ESAYFFFEKN LKDVSFRLGS FNLEKVENPA
EVIRELIAYA LDTIAENQAK NEHLQKENER LLRDWNDVQG RFEK
pSIM8
pSIM33
pSIM123
NMR intensity loss
Unassigned prolines
Others unassigned/ overlapping
1H-15N BEST-TROSY spectra of XRCC4 1-164 +/- mSUMO2
R3
M59
D57
V158
V104
H143
K72
A73
Chemical shift δ1H (ppm)
Chemical shift δ15N (ppm)
E98
Q145
E25
K26
N148
L144
F106
E49
K99
K4
Q138
K102
B
15N-XRCC4 1-164 + diSUMO2
C
15N-XRCC4 1-164 + 4xSUMO2
D
15N-XRCC4 1-164 + mSUMO1
90°
E
XRCC4 head domain
Electrostatic surface potential
Orientation
- 5
*kT/e
5
SUMO2-binding surface

### Figure S5

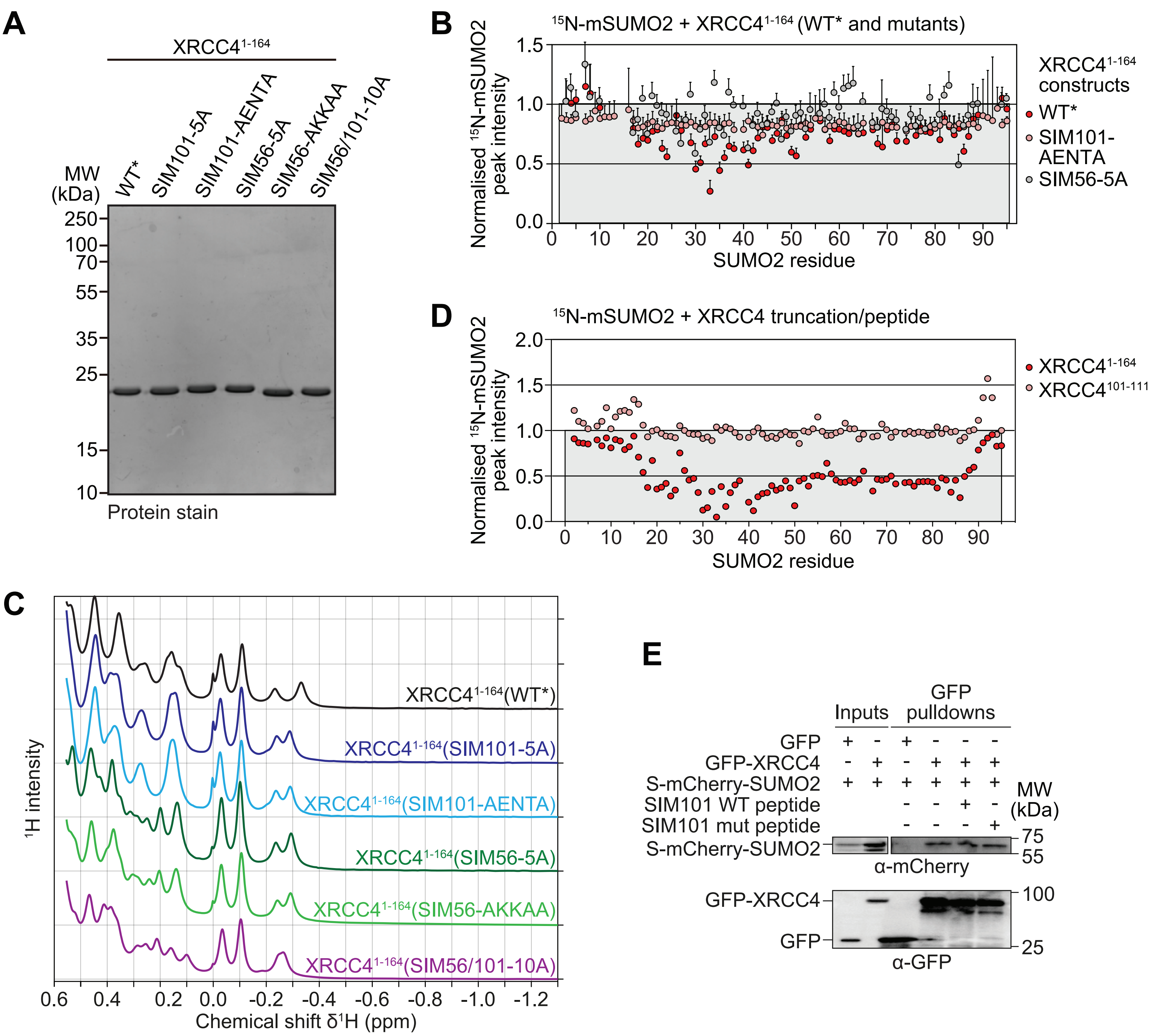

A
XRCC4 1-164
WT*
SIM101-5A
SIM101-AENTA
SIM56-5A
SIM56-AKKAA
SIM56/101-10A
MW (kDa)
250
100
70
55
35
25
15
10
Protein stain
B
15N-mSUMO2 + XRCC4 1-164 (WT* and mutants)
Normalised 15N-mSUMO2 peak intensity
SUMO2 residue
XRCC4 1-164 constructs
WT*
SIM101-AENTA
SIM56-5A
C
XRCC4 1-164(WT*)
XRCC4 1-164(SIM101-5A)
XRCC4 1-164(SIM101-AENTA)
XRCC4 1-164(SIM56-5A)
XRCC4 1-164(SIM56-AKKAA)
XRCC4 1-164(SIM56/101-10A)
1H intensity
Chemical shift δ1H (ppm)
D
15N-mSUMO2 + XRCC4 truncation/peptide
Normalised 15N-mSUMO2 peak intensity
SUMO2 residue
XRCC4 1-164
XRCC4 101-111
E
Inputs
GFP pulldowns
GFP
GFP-XRCC4
S-mCherry-SUMO2
SIM101 WT peptide
SIM101 mut peptide
MW (kDa)
75
55
α-mCherry
GFP-XRCC4
GFP
100
25
α-GFP

### Figure S6

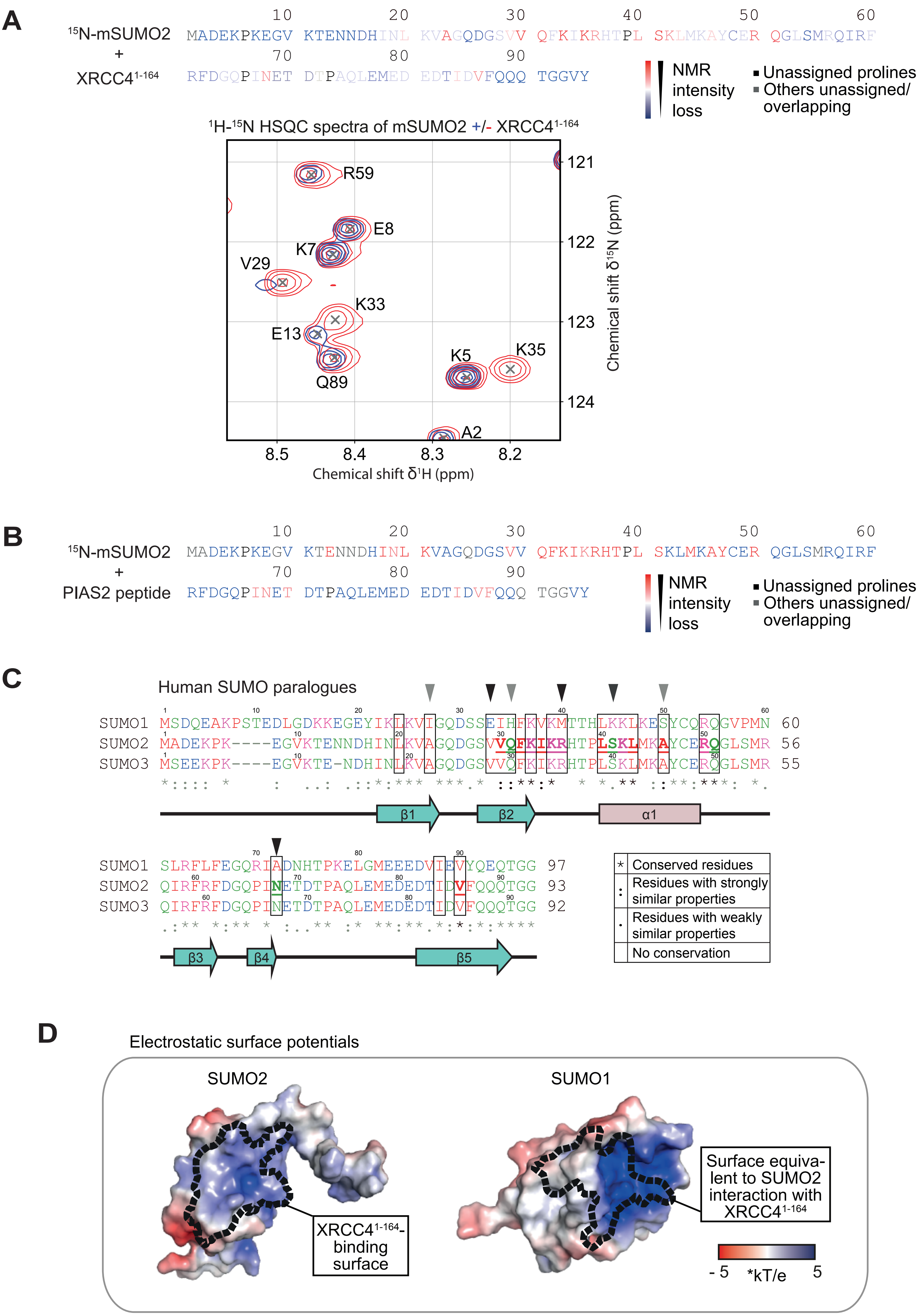

A
15N-mSUMO2
+
XRCC4 1-164
10 20 30 40 50 60
MADEKPKEGV KTENNDHINL KVAGQDGSVV QFKIKRHTPL SKLMKAYCER QGLSMRQIRF
70 80 90
RFDGQPINET DTPAQLEMED EDTIDVFQQQ TGGVY
NMR intensity loss
Unassigned prolines
Others unassigned/ overlapping
1H-15N HSQC spectra of mSUMO2 +/- XRCC4 1-164
R59
E8
K7
V29
K33
E13
Q89
K5
K35
A2
121
122
123
124
8.5
8.4
8.3
8.2
Chemical shift δ1H (ppm)
Chemical shift δ15N (ppm)
B
15N-mSUMO2
+
PIAS2 peptide
10 20 30 40 50 60
MADEKPKEGV KTENNDHINL KVAGQDGSVV QFKIKRHTPL SKLMKAYCER QGLSMRQIRF
70 80 90
RFDGQPINET DTPAQLEMED EDTIDVFQQQ TGGVY
NMR intensity loss
Unassigned prolines
Others unassigned/ overlapping
C
Human SUMO paralogues
SUMO1 MSDQEAKPSTEDLGDKKEGEYIKLKVIGQDSSEIHFKVKMTTHLKKLKESYCQRQGVPMN 60
SUMO2 MADEKPK----EGVKTENNDHINLKVAGQDGSVVQFKIKRHTPLSKLMKAYCERQGLSMR 56
SUMO3 MSEEKPK----EGVKTE-NDHINLKVAGQDGSVVQFKIKRHTPLSKLMKAYCERQGLSMR 55
β1
β2
α1
SUMO1 SLRFLFEGQRIADNHTPKELGMEEEDVIEVYQEQTGG 97
SUMO2 QIRFRFDGQPINETDTPAQLEMEDEDTIDVFQQQTGG 93
SUMO3 QIRFRFDGQPINETDTPAQLEMEDEDTIDVFQQQTGG 92
β3
β4
β5
* Conserved residues
: Residues with strongly similar properties
. Residues with weakly similar properties
No conservation
D
Electrostatic surface potentials
SUMO2
SUMO1
XRCC4 1-164-binding surface
Surface equivalent to SUMO2 interaction with XRCC4 1-164
- 5
*kT/e
5

### Figure S7

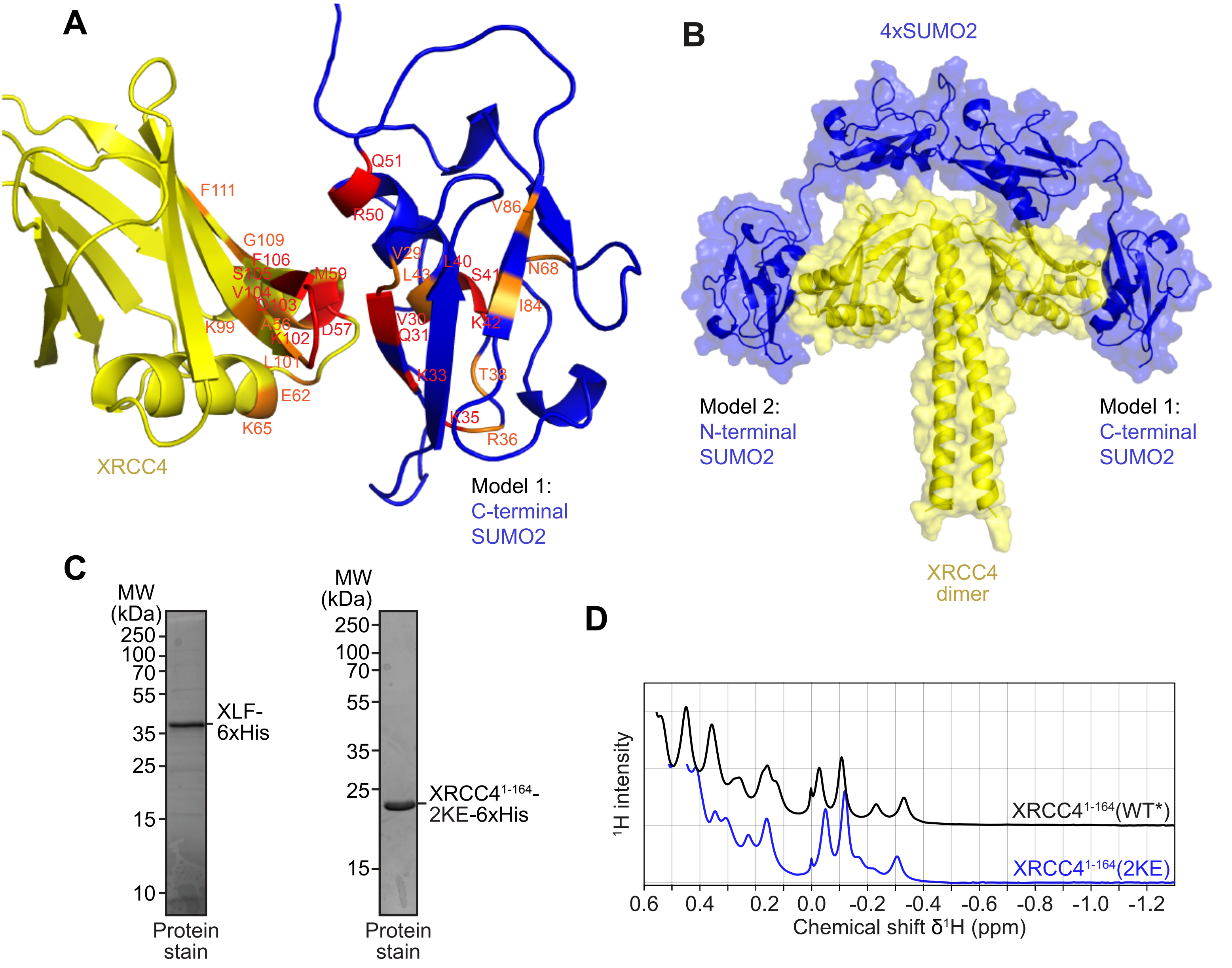
A
F111
G109
F106
S105
V104
D103
M59
K99
A56
K102
D57
L101
E62
K65
XRCC4
Q51
R50
V86
V29
L40
L43
S41
N68
V30
Q31
K42
I84
K33
T38
K35
R36
Model 1:
C-terminal
SUMO2
B
4xSUMO2
Model 2:
N-terminal
SUMO2
Model 1:
C-terminal
SUMO2
XRCC4
dimer
C
MW
(kDa)
250
100
70
55
35
25
15
10
XLF-
6xHis
Protein
stain
MW
(kDa)
250
100
70
55
35
25
15
XRCC4 1-164-
2KE-6xHis
Protein
stain
D
¹H intensity
XRCC4 1-164(WT*)
XRCC4 1-164(2KE)
0.6
0.4
0.2
0.0
-0.2
-0.4
-0.6
-0.8
-1.0
-1.2
Chemical shift δ¹H (ppm)

### Figure S8

**A**

| | | | | | | |
|---|---|---|---|---|---|---|
| | 10 | 20 | 30 | 40 | 50 | 60 |
| $^{15}$N-mSUMO2 | MADEKPKEGV | KTENNDHINL | KVAGQDGSVV | QFKIKRHTPL | SKLMKAYCER | QGLSMRQIRF |
| + | 70 | 80 | 90 | | | |
| XRCC4$^{FL}$ | RFDGQPINET | DTPAQLEMED | EDTIDVFQQQ | TGGVY | | |

NMR intensity loss

■ Unassigned prolines
■ Others unassigned/ overlapping
